# T cell degranulation and payload delivery are modulated in space and time

**DOI:** 10.64898/2026.07.23.740126

**Authors:** Kevin L. Scrudders, Geoffrey H. Graff, Laurent Limozin, Kheya Sengupta, Shalini T. Low-Nam

## Abstract

Release of destructive payloads through degranulation is a hallmark of T cell cytotoxicity. Upon stimulation, tens to hundreds of granules are rapidly delivered to the interface with a target cell but only a small number of degranulation events occur. Two forms of degranulation have been observed and differ in the physical form of the payload released. The deposition of insoluble particles encapsulated in a proteinaceous shell may be more potent than release of purely soluble factors. The selection mechanisms and triggers for degranulation are still poorly understood, partly due to the lack of tools to dynamically map and distinguish each mode. Here we use high spatiotemporal resolution *in vitro* assays, coupled to a novel label-free strategy to characterize and track released particles, to show that degranulation is regulated not only by chemical but also physical aspects of the activation inputs. Furthermore, the released, insoluble particles are transported within the intermembrane junction with a preferred directionality. Immobilized stimulatory ligands favored the soluble degranulation mode suggesting that the particulate release is pertinent to mobile, physiological immune interfaces. Engineered chimeric antigen receptor (CAR) T cells have a defect in centralizing released particles, which may explain dampened toxicity and off-target tissue damage. Our results demonstrate how degranulation events are likely regulated by a combination of physical, chemical, and signaling mechanisms. We anticipate these findings will generate new strategies to bias degranulation outcomes and optimize the destructive capacity of single cytotoxic T cells. In the case of engineered T cells, efforts to restore the normal transport of insoluble particles could contribute to stronger therapeutics for cellular immunotherapy.

---

T cells generate cytotoxicity by releasing perforin and granzyme molecules into the intermembrane junction with a target cell and inducing cell death in the latter^1^. Complex inputs, integrated at the interface, may culminate in the polarized trafficking of tens to hundreds of lytic granules and degranulation of cytotoxic contents derived from lysosomal compartments^2,3^. Recent work proposes distinct modes of degranulation involving either granules that release purely soluble contents or granules composed predominantly of insoluble material^4–6^. The insoluble particles, which have been called supramolecular attack particles (SMAPs), encapsulate perforin, granzyme, and serglycin in a glycoprotein shell comprised largely of thrombospondins^5–7^. These particles have been shown to generate potent, autonomous cytotoxicity^7^. The molecular thresholds that control and select between these exocytic modes are of great interest to control cytotoxic outcomes but remain unclear. How membrane sites for degranulation are licensed is also notable when considering that degranulation is a rare event with an estimated 2-12 granules released following activation^8–10^. The preservation of the majority of the payload could contribute to serial killing of multiple targets^11,12^.

The spatial and temporal scales of degranulation contribute to the complexity of its detection. Visualization of cytotoxic granules and membrane fusion sites by electron microscopy and fluorescence imaging has set the size scale of hundreds of nanometers in diameter per granule and topographic features for exocytic release^8,9,13–17^. Live cell imaging has also demonstrated that the lifetime of degranulation events can vary over an order of magnitude in time^10,18^. Membrane fusion can occur within hundreds of milliseconds while the full lateral diffusion of the granule membrane to mix with the adjacent plasma membrane may take up to seconds. Much of the signaling and fusion machinery has been characterized and points to calcium-dependent mobilization of granules that are trafficked along microtubule and actin fibers to the intermembrane junction. There is strong evidence that the adhesion characteristics of the intermembrane junction provide a checkpoint for granule fusion^2^. Thus, the interplay of mechanics and biochemical regulation shapes the degranulation profile.

Native T cell receptors (TCR) sensitively trigger in response to peptide-loaded major histocompatibility complexes (pMHCs). Abundant evidence demonstrates that robust responses, including cytotoxicity, are generated following small handfuls of binding interactions, collected stochastically^19–24^. The spatial and temporal correlation of binding sequences sets the activation threshold by generating local signaling assemblies that promote signal amplification for early decision-making outcomes^19,20^. Of additional interest is whether these mechanisms are engaged when T cells are activated through orthogonal machinery, such as artificial receptors. Engineered T cells based on chimeric antigen receptors (CARs) coopt the T cell signaling machinery to repurpose cytotoxicity toward destruction of tumor and autoimmune targets^25,26^. However, substantial evidence suggests that CARs weakly propagate signals and kill less efficiently than native T cells, raising questions about how different degranulation modes are mobilized and if these processes are tunable^27,28^.

There is a need to bridge electron microscopy characterization of granules and membrane features at fusion sites and live cell fluorescence imaging of the release of granule contents^2,5,8,10,29^. Here, we address this gap by using high resolution, surface-selective imaging to simultaneously monitor degranulation of polarized granules and the membrane landscape in primary human T cells. Cells were exposed to either substrates displaying immobile adhesion and stimulatory ligands or ligands displayed on mobile supported lipid bilayer (SLB) platforms which are known to differentially impact cell spreading and activation^30,31^. In each case, cells were capable of generating both degranulation modes. In response to immobilized ligands, there was a weighting toward release of granules containing soluble contents, suggesting that the reorganization of bound receptors at the interface is a component of particle release. Release of soluble contents typically occurred first and toward the periphery of the contact zone. The weighting of degranulation modes toward release of insoluble particles on mobile surfaces prompted questions about the role of signaling in biasing the outcome. Using a second generation chimeric antigen receptor (CAR) T cell model, we found that there is a defect in transporting released particles toward the center of the interface, presumably mapping to the weaker signaling ascribed to these cells^27,32^. These data demonstrate that T cells integrate physical and biochemical mechanisms to effect degranulation and control potency toward target cells.

## Results

### Primary T cell degranulation on glass substrates presenting immobilized ligands

Considerable evidence suggests that submaximal degranulation of cytotoxic payloads is a characteristic of immune cell degranulation. Furthermore, small numbers of events resulting in target destruction^33^ emphasizes the need to understand how cells control these exocytic mechanisms. Detection of individual degranulation events required discrimination of granule fusion from the ensemble of granules that were polarized to the T cell surface following stimulation. Primary human T cell lytic granules were labeled with membrane-permeable LysoTracker, a pH-sensitive fluorescent dye that accumulates within acidic compartments. We used anti-CD3 and anti-ICAM-1 antibody-coated glass to strongly trigger TCRs and integrins, respectively, and generate robust polarization and degranulation responses (Fig. 1A). An interleaved imaging strategy maximized the detection of degranulation events and corresponding membrane geometries. Label-free interference reflection microscopy (IRM), at low frequency, was used to detect the interfacial contact area and features of the PM topography (Fig. 1B; Fig. S1)^34,35^. Cells were observed from their initial contact and landing to capture the full sequence of polarization and degranulation events (Fig. 1B; Movie S1). LysoTracker fluorescence was monitored using total internal reflection fluorescence (TIRF) microscopy at a typically rapid, streaming rate of 10 Hz (Fig. S2). Ensemble polarization was assessed based on aligning cells to the time of landing and measuring the accumulation of granules at the T cell surface (Movie S1). The extent of polarization was higher on those surfaces that triggered the TCR, consistent with a stronger activation (Fig. 1C). ELISA-based detection of granzyme B in supernatants collected 30 minutes after T cell exposure to the solid substrates showed the expected high levels following activation on these surfaces (Fig. S3).

**Figure 1.**
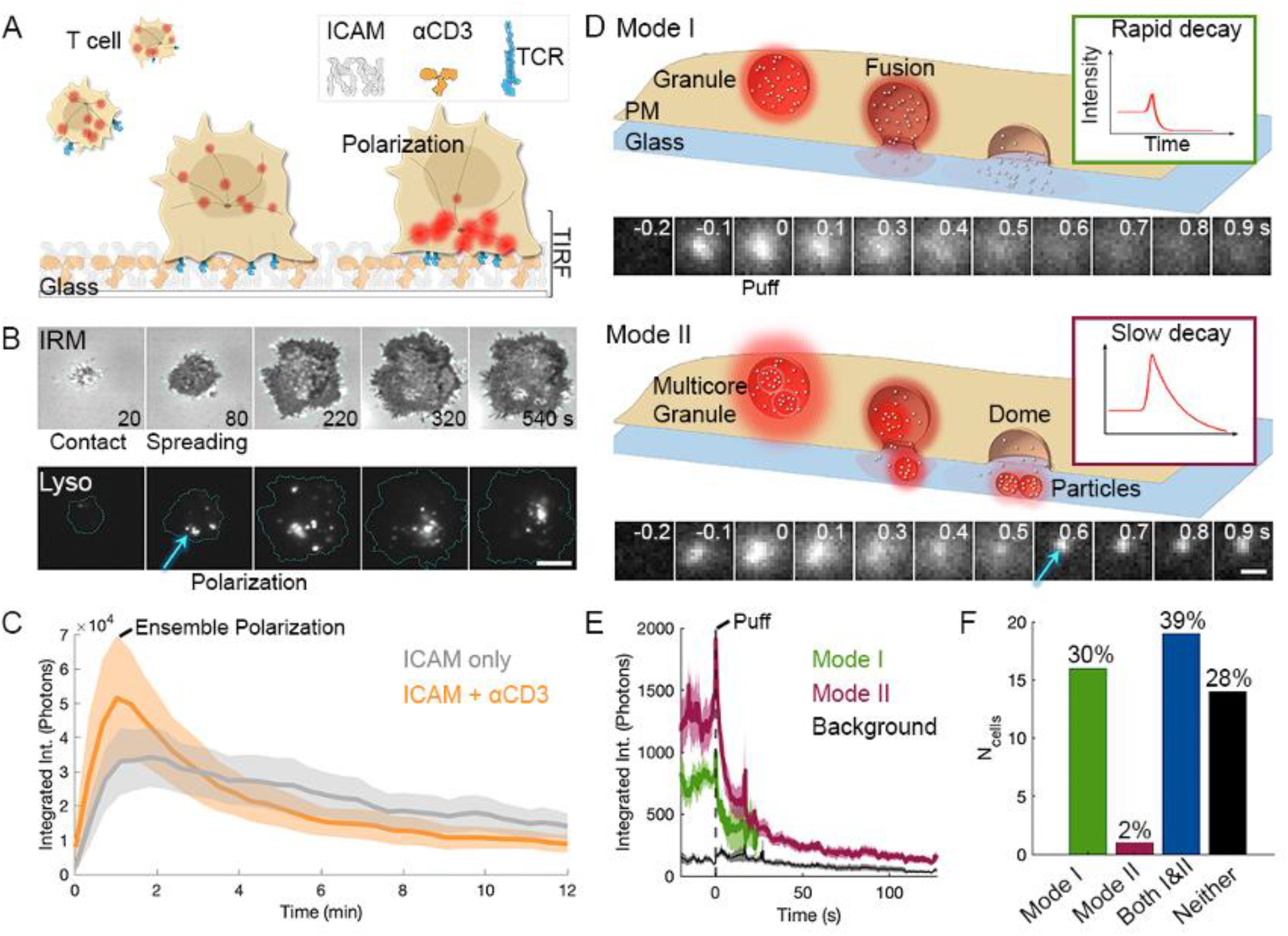
Robust T cell polarization and two modes of degranulation observed on glass surfaces. **A**. Schematic of primary human T cell landing on poly-D-lysine-coated glass substrates derivatized with ICAM-1 (ICAM) and anti-CD3. Cells are loaded with LysoTracker, enabling visualization of polarization in a total internal reflection fluorescence (TIRF) configuration. **B**. T cell landing, spreading, and polarization (arrow) of LysoTracker (Lyso)-labeled granules visualized using interference reflection microscopy (IRM) and TIRF. Scale bar = 5 microns. **C**. Quantification of ensemble polarization of T cells on resting (ICAM-1 only; gray) or activating (ICAM-1+anti-CD3; orange) surfaces. Individual traces of fluorescence intensity were aligned to cell landing. Average intensity is shown as a solid line with shaded areas reflecting the standard deviation. Each trace includes >40 cells across >4 substrates and >2 experiments. **D**. Schematics for Mode I and Mode II degranulation showing LysoTracker-labeled granules and particles, fusion, and release of material. Insets show the corresponding fluorescence intensity decay profiles. Below each schematic is a raw data example of a degranulation event showing the fluorescent puff at fusion (time = 0 s). Arrow in Mode II points to the particulate material that was released. **E**. Ensemble average of Mode I (186 events) and Mode II (55 events) degranulation events sorted based on the decay time (tau). Background intensity trace shown in black. For each trace, the standard deviation is shown as a lightly shaded area around the average. Percentages indicate the fraction of all events for each mode. **F**. Number of cells observed to land and undergo Mode I-only, Mode II-only, both modes, or neither form of degranulation, respectively. Percentage of all cells is show above each corresponding bar.

The different modes of degranulation would predict distinct fluorescence signatures as a function of time (Fig. 1D). We define Mode I granules to release soluble perforin and granzyme contents, whereas Mode II granules deposit insoluble material onto the underlying substrate, in addition to releasing some soluble contents. By using rapid acquisitions, the behavior of individual granules at the surface could be tracked and assessed. Upon fusion with the plasma membrane, the pH of granule lumen was rapidly neutralized, leading to a quenching of LysoTracker fluorescence. Thus, granule fusion was marked by a transient, sharp peak in intensity (exocytic ‘puff’) followed by rapid fluorescence decay and the radial diffusion of the signal (Fig. 1D). An example Mode I puff event shows the lateral redistribution of the fluorescence signal whose signal is quickly lost (Fig. 1D; Movie S2). Mode II events showed a persistent fluorescent spot after the puff that could be observed up to tens of seconds and was characteristic of LysoTracker signal inside the particle shell and, thus, shielded from fluorescent quenching (Fig. 1D; Movie S3).

Given the potent stimulation on the glass surface, hundreds of individual degranulation events were captured over tens of cells. Capturing cell landing ensured maximal detection of degranulation starting from the earliest signal integration and without the requirement for formation of a mature immunological synapse. Quantitative features of the loaded granules before and after fusion were extracted and aligned to the puff as a proxy for the time of degranulation. The fluorescence decay rates after fusion were well fit by a single exponential. These signatures permitted parsing into two groups (Mode I tau = 1.98 s; Mode II tau = 22.92 s), consistent with the two degranulation modes and similar to prior reports^36^ (Fig. 1D-1E). Most cells displayed both modes of degranulation and only rarely was Mode II degranulation the exclusive mechanism of material release, suggesting that fusion of these granules alone is disfavored and may require some of priming of the cellular machinery by Mode I events (Fig. 1F). Also notable was that even on these highly triggering surfaces, some cells did not undergo degranulation (Fig. 1F). Therefore, a subset of the T cell population is predominating the cytotoxic response, similar to some other observations, including clinical results using engineered T cells^37–39^.

### Mode II events generate a striking membrane signature

The substantial membrane deformation that is expected to accompany degranulation^40,41^ events prompted us to measure if the inherent topographic information embedded within interference reflection microscopy (IRM) images (Fig. 1B) could be used to distinguish the degranulation modes. Despite the rapidity of the changes in the fluorescent signature of LysoTracker, the timescale of the membrane remodeling during exocytosis could be slower, such as predicted by models in neurons^42^ and in NK cells^43^. IRM provides label-free information on membrane topography^34,35^. An IRM image is formed by interference of light rays reflected at two or more interfaces. We frequently observed a strong correspondence between bright LysoTracker-positive granules near the surface and distinct IRM features, most often a dark black spot surrounded by a white halo (Fig. 2A). This pattern covered several pixels (>4 pixel radius) and took on a circular symmetry reminiscent of a spherical object. This correlation between channels was most often observed for long-lived fluorescent objects, leading us to predict they were related to Mode II events. To explore these signatures in greater detail, we interleaved, instead, high frequency (10 Hz) IRM imaging with infrequent (0.1 Hz) imaging of LysoTracker-labeled granules. In this case, the exact timing of degranulation could not be confirmed given the slower fluorescent imaging. An example shown in Figure 2B shows a long-lived granule whose appearance corresponded to a striking IRM feature that persisted for many tens of seconds, even after the fluorescent signal photobleached (Fig. 2B). This sustained signal in IRM was consistent with released particles that were deposited onto the underlying glass surface. To confirm release of granules and deposition, we concentrated on datasets in which T cells were observed to retract their membranes after landing and spreading on the activating surface (Fig. 2C, Movie S1). As expected, we, again, observed correspondences between LysoTracker-labeled granules and IRM signatures. When these events occurred near the edge of the contact area and the T cell retracted, the presumptive Mode II particle was observed to be left behind on the glass and could be discriminated as a dark IRM object and a spherical one in diascopic imaging (Fig. 2C and Fig. S4).

**Figure 2.**
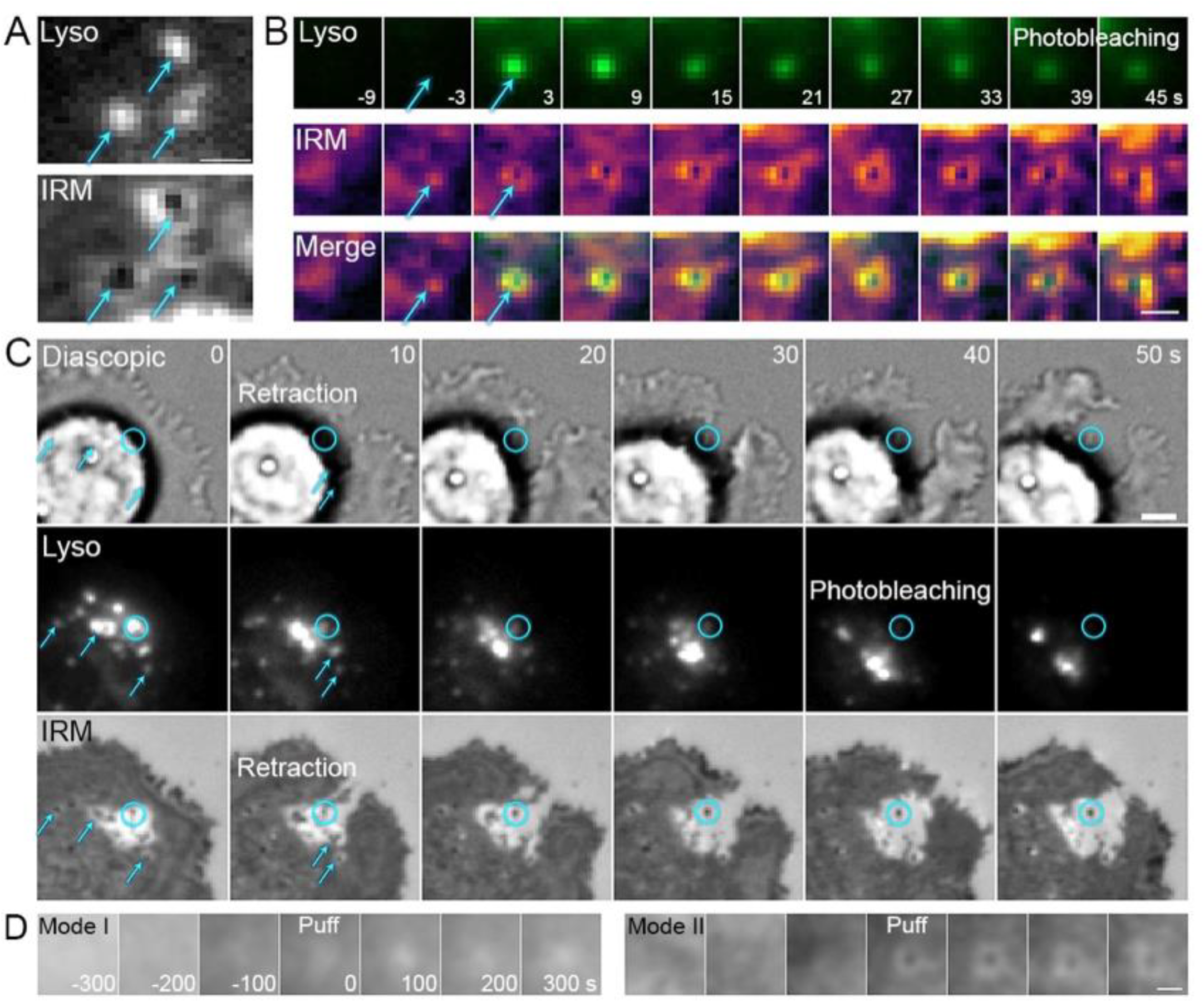
Mode II events have striking membrane signatures and deposit material onto the underlying substrate. **A**. Images of LysoTracker (Lyso)-labeled polarized granules visualized in TIRF and corresponding features in Interference Reflection Microscopy (IRM) image. Arrows indicate individual granules in each channel. Scale bar = 2 microns. **B**. Montages of a Mode II event imaged with rapid IRM (10 Hz; middle row) and slow LysoTracker (0.167 Hz; top row). Overlay shown in bottom row. Fusion event (puff in fluorescence; 0 s) was not directly visualized. Timestamps show the presumed times of the event. Arrow at -3 seconds indicates a feature in IRM that preceded the expected puff frame. Arrow at 3 seconds shows the strong spatial colocalization of the LysoTracker and IRM signals. Scale bar = 1 micron. **C**. Montages of a T cell that underwent polarization and part of the membrane retracted (starting at time = 10 s). Diascopic (top row), LysoTracker (middle row), and IRM (bottom row) shown. Arrows indicate example features at corresponding positions in both Lyso and IRM channels. Circle indicates an example feature that corresponds between all 3 channels and was observed to remain on the glass after T cell retraction. Scale bar = 2 microns. **D**. Ensemble average montages of Mode I (Left) and Mode II (Right) events using classifications from fluorescent traces (see Figure 1E). Scale bar = 1 micron.

To confirm if IRM could be used to monitor both modes of degranulation, we used the classification of events from fluorescence (Fig. 1E), in which degranulation was observed, to inspect the corresponding, low temporal resolution IRM images (Fig. S5). Averaging over events aligned to the puff showed that Mode I events tended to have a subtle signature in IRM (Fig. 2D). By comparison, the Mode II event ensemble average had the dark spot and white halo, with that signature being persistent over many tens of seconds (Fig. 2D). We interpreted this to mean that IRM could be used to monitor the dynamics of Mode II degranulation events without a requirement to directly observe the fluorescent puff.

### Biophysical features of cytotoxic particle release

The qualitative observations of coincident lytic granule and membrane features prompted a more detailed characterization of the geometric and optical features that were embedded in the IRM data. We suspected that Mode II particles generated the distinct spot and halo IRM pattern due to differences in refractive index and consistent with dense, proteinaceous cores expected especially for SMAPs^6^. From the literature, the diameter of cytotoxic granules is expected to be in the range of 100 to 1000 nanometers. So far, the formalism of IRM or the closely related RICM^31,35,44–46^ was usually applied to much larger particles (at least 5 microns)^47^. Very small objects are expected to scatter light, instead of reflecting it^48^. However, the typical, expected size of granules considered here is considerably larger than the limit of Rayleigh scattering usually used to quantify interference scattering. We therefore decided to exploit conventional RICM analysis to simulate the image of objects of sizes comparable to Mode II granules. The modeling was parsimonious and focused on the minimal biological and optical parameters that were expected to characterize degranulation events.

To start, Mode I and Mode II events before and after fusion were considered with simplistic geometries (Fig. 3A). The membrane was modeled as a dome-shape (concave hemisphere) contiguous with the cell membrane and approximated as step-wise parallel planes. In the case of Mode II degranulation, the particles were modeled as a vertical cylinder with height and base diameter directly under the membrane-dome (Fig. 3A). For both experimental and simulated data, the background, corresponding to reflection from glass, was set to 1 and the rest of the image was normalized accordingly. While there was not a quantitative match between intensities, the overall pattern matched qualitatively very well (Fig. 3A, Fig. S6). Mode I granules, estimated to be 200 nm in diameter, showed very little contrast in the simulations. This compared well with raw data (Fig. 3A) of single events. The white spot seen after fusion in the simulation compared favorably with the ensemble average of many events (Fig. 2D). This suggests that, over many Mode I events, aligned to the time of degranulation, a subtle membrane signature is evident. Overall, given the absence of striking features in typical data imaging IRM at low frequency, we interpret this to mean that the Mode I granule membrane fuses and becomes contiguous with the plasma membrane, mostly adopting its topography, within 10 seconds.

**Figure 3.**
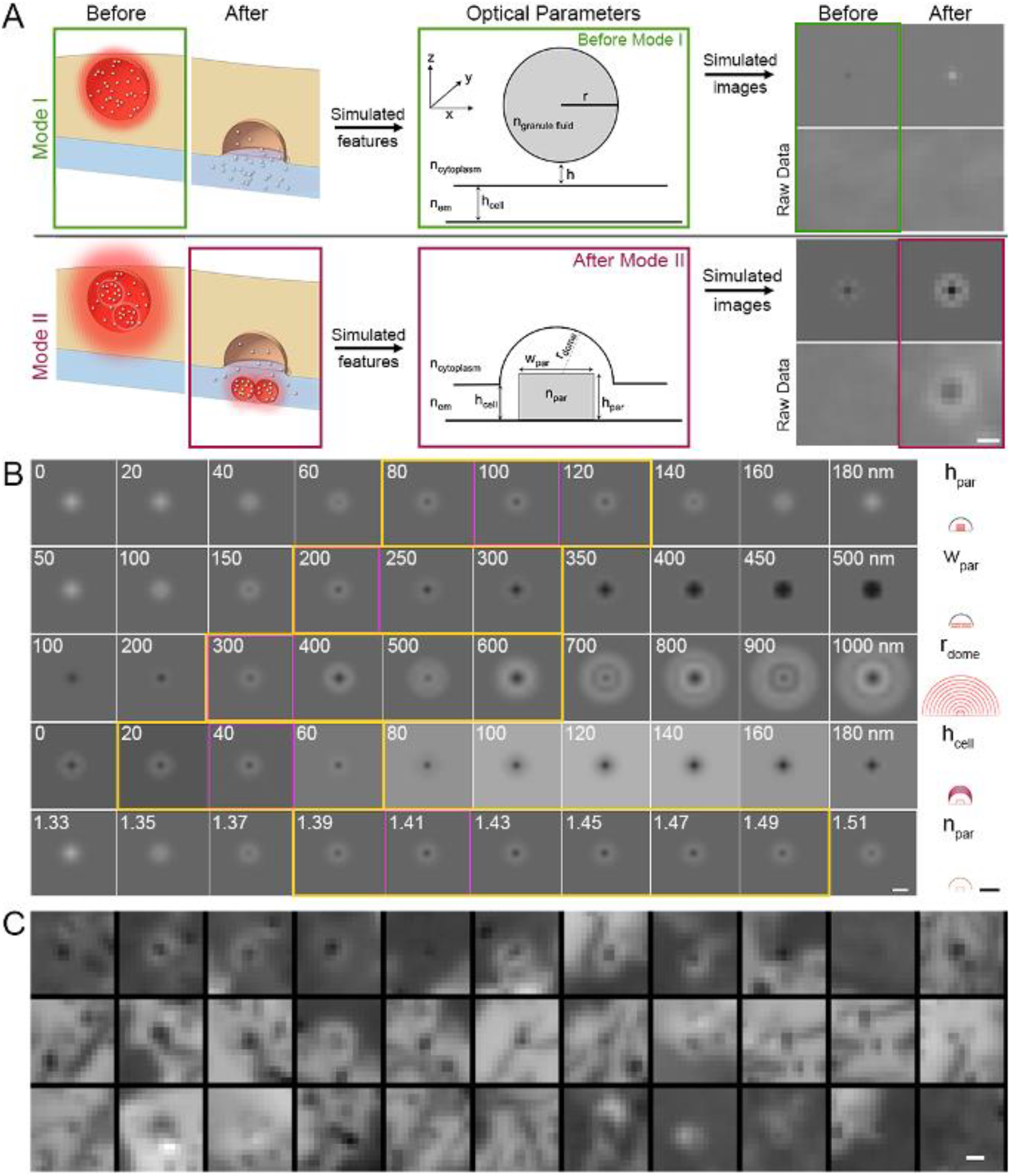
Simulation of IRM signatures for degranulation events, matched to raw data, provide features especially of Mode II events. **A**. Schematic of Mode I and Mode II degranulation events that are simulated to generate expected IRM signatures. Left: Before and after features of Mode I and Mode II degranulation. Center: Corresponding components of IRM simulations. Center, top is Mode I before degranulation. Center, bottom is after Mode II degranulation. Abbreviations: n_granule fluid_ is index of refraction of granule contents; n_cytoplasm_ is index of refraction of cytoplasm; n_em_ is index of refraction of extracellular medium; r is granule radius; h is height above membrane; h_cell_ is membrane height above substrate; r_dome_ is radius of membrane dome; w_par_ is width of particle; h_par_ is height of particle; n_par_ is index of refraction of particle. Right: Simulated and raw data images corresponding to the before and after cases for each degranulation mode. Scale bar is 0.5 microns. Simulation parameters are provided in the Materials and Methods. **B**. Simulated features of Mode II IRM signatures for a range of parameters. Each row corresponds to the parameter that was independently varied, as shown to the right. The feature is shown on the far right using the same scheme as in (A) center column. Magenta boxes indicate the parameters that returned simulations most similar to the raw data. The range of data boxed in yellow indicate the range of values that bore similarities to the raw data. Scale bar is 0.5 microns. **C**. Ensemble average images of Mode II events showing the average of many frames acquired by IRM streaming at 10 Hz. Scale bar is 0.5 microns.

For Mode II degranulation, the simulated and experimental images were qualitatively similar. For sequences in which IRM was sampled at a fast rate, the approach of the granule provided a signature that looked similar to the simulation (Fig. 2B and Fig. 3A). We further simulated the Mode II granules over an extended range of parameter values to determine which sets of features most closely resemble the IRM data from experiments (Fig. 3B; Fig. S6). The deposited particles had the most striking features in both the raw and simulated data with the characteristic pattern of a dark spot surrounded by a bright halo. The simulations put a strong constraint on particle size which we estimate to be about 100 nanometers in height with a width of approximately 200 nanometers. These estimates match measurements from cryo-electron microscopy and super-resolution imaging^6^. The radius of the domed, overlying membrane was estimated to be between 300-600 nanometers and 40 nanometers above the substrate, at the lowest point of the dome. The height of the plasma membrane closely matches the estimated distance of adhesion interactions^49^, suggesting high fidelity in the simulations. The index of refraction of the particles falls into the range of 1.39-1.49, which is comparable to other protein-dense organelles (Fig. 3B). Collectively, the simulations generate signatures that closely resemble the average feature from Mode II events sampled using fast IRM imaging (Fig. 3C, Movie S4-S9).

### Spatiotemporal features of released Mode II particles

The spatial and temporal resolution achieved in these measurements enabled us to interrogate where and when each degranulation event occurred. The sequence of degranulation events for each cell after landing on the glass surface were plotted (Fig. 4A). Mode I degranulation occurred first ∼80% of the time (Fig. 4B). This suggests that these granules are either trafficked to the interface first or that the assembly of fusion machinery and actin depletion necessary for fusion favor these events, perhaps filtering based on the size of the granule. On the contrary, the last event observed, typically around 4-5 minutes after landing, was equally likely to be of the Mode I or Mode II variety (Fig. 4C). These data argue that degranulation modes and timing may be ordered, especially at the onset of degranulation. Further, recent data reiterating a role for cytoskeletally-docked ICAM in creating immobile sites for adhesion suggest that Mode I events occurring early and toward the edges of the contact zone may be physiologically-relevant^50^.

**Figure 4.**
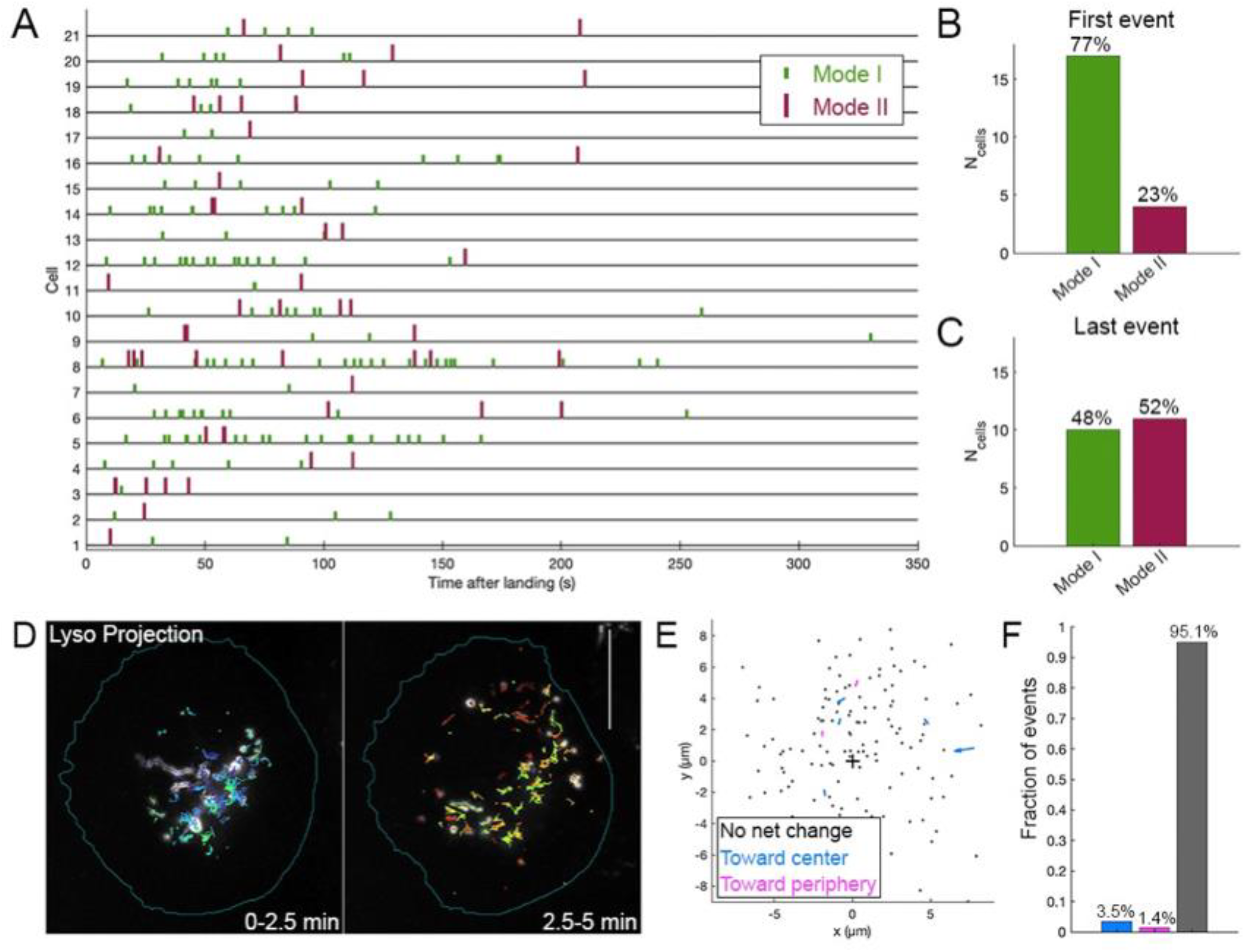
Degranulation events mapped onto the interface show Mode I occurs earlier than Mode II and Mode II degranulation on glass substrates results in immobile particles. **A**. Event timelines for cells with at least one event each of Mode I and Mode II. Cells are aligned to time of landing (t = 0). **B**. Fraction of first degranulation events per cell of each mode. **C**. Fraction of last degranulation events per cell of each mode. **D**. Tracked granule trajectories plotted on maximum projection for an example cell on glass. LysoTracker data are projected over 2.5-minute intervals and each tracked granule longer than 500 ms are plotted in unique colors. Scale bar = 10 microns. **E**. Displacement vectors for Mode II degranulation events. All events shown across 33 cells where the center of each cell interface is assigned the coordinate (0,0). **F**. Summary of data in (E) showing the fraction of events for which the Mode II particle moves toward or away from the center, or has no negligible displacement, respectively.

Tracking of LysoTracker-labeled granules permitted characterization of preferred sites for degranulation (Fig. 4D). The onset of polarization included many granules near the center of the contact zone, but Mode I events were frequently localized to the periphery of the contact zone (Fig. 4D; Fig. S7). This suggests that fusion mechanisms are also spatially regulated. Mode II degranulation occurred closer to the center of the contact zone (Fig. S7). This organization suggests that the released material may be focused within the synapse to generate a high effective concentration and may minimize lateral diffusion away from the target cell. The movement of released particles, based on tracking the IRM feature was quantified to determine the net displacement and its directionality. On the glass surface, the particles were mostly immobile, consistent with some sticking to the underlying, coated substrate (Fig. 4E-4F).

### Degranulation behaviors within a mobile, complex intermembrane junction

To further explore the two degranulation modes in the context of a true intermembrane junction, we replaced rigid glass substrates with supported lipid bilayers (SLBs) functionalized with mobile ligands for TCR triggering (Fig. 5A). The bilayer system mimics a mobile target cell membrane, with diffusing ligands, while maintaining optical accessibility^51–53^. SLBs presenting a high density of ICAM and anti-CD3 antibodies at moderate densities were used as mimics of antigen presenting cells. Just like the observations on glass, LysoTracker-loaded primary human T cells were recorded to observe puff events (Movie S10) from the onset of landing. One substantial difference compared to degranulation on glass was a higher weighting to Mode II events. On the SLB, Mode I and Mode II events were, approximately, evenly split (Fig. 5B). Thus, the mobility of ligands for TCR triggering are implicated in the outcome of the degranulation program.

**Figure 5.**
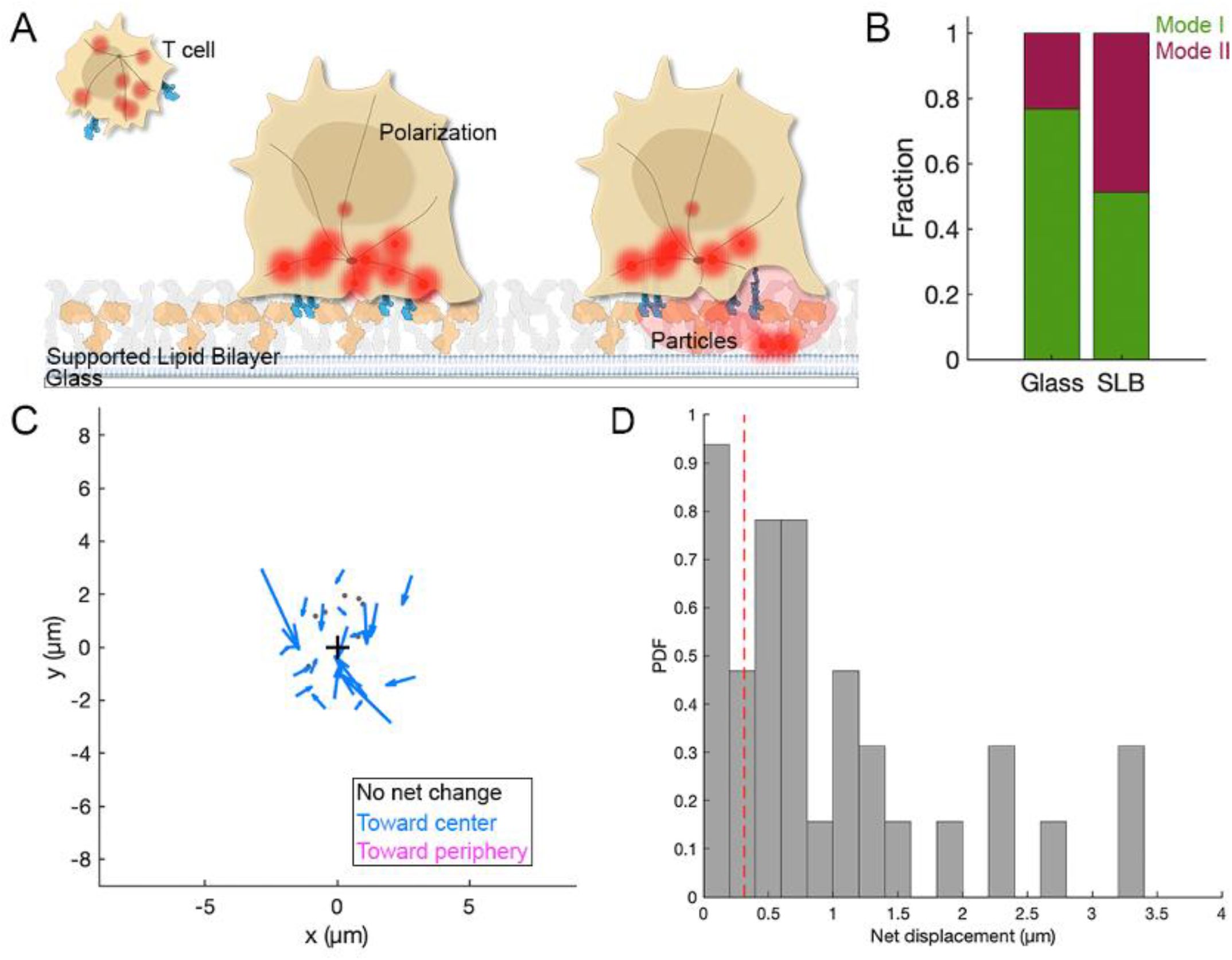
Mobile TCR ligands shift the ratio of degranulation events and Mode II released particles are transported to the center of the interface. **A**. Schematic of primary human T cell activation on supported lipid bilayers (SLBs) presenting mobile ICAM and anti-CD3 ligands. Cells were loaded with LysoTracker and exposed to activating substrates to promote polarization and degranulation. **B**. Fraction of degranulation events of Mode I and Mode II on each of glass (immobile ligand) and SLB (mobile ligand) surfaces. **C**. Displacement vectors for Mode II degranulation events. All events shown across 11 cells where the center of the interface is assigned the coordinate (0,0). **D**. Histogram of Mode II particle displacement. Red, dashed line indicates cutoff for immobile particles (<0.3 micron displacement).

The particles released on SLBs were observed to undergo movement in IRM, predominantly toward the center of the intermembrane interface (Fig. 5C). Tracks of granules marked by IRM features alongside the pre- and post-puff LysoTracker signals showed correlated motion. Within the scope of these measurements, we did not observe any particles to undergo excursion toward the periphery. This suggests that sorting of molecules during maturation of the immunological synapse (IS)^53,54^ imposes transport on the particles (Fig. S8). Presumably, the movement of TCRs toward the central supramolecular activation cluster (cSMAC) is key to this process. Some of the particles experienced net displacements of more than 1 micron (Fig. 5D), consistent with measurements of TCR net transport during IS formation^55^. These data are consistent with our hypothesis that T cells enhance cytotoxic function by increasing the destructive capacity at the center of the interface.

### Engineered T cell degranulation has an altered balance of degranulation modes

The differences that have been noted in TCR and chimeric antigen receptor (CAR) T cell signaling^32,56^ and effector function prompted us to question how degranulation differs during TCR-independent triggering (Fig. 6A). The damped signaling by CARs has been the focus of ongoing engineering efforts to strengthen activation outcomes. We focused on a universal CAR T cell that expresses an anti-FITC CAR (Fig. 6A). To promote engagement of this anti-FITC CAR T cell with a target cell, we used a folate-fluorescein bispecific adapter, termed EC-17 (Cytalux^™^), to form a bridge between the anti-FITC CAR on the T cell and a folate receptor on a cancer cell (Fig. 6A)^57,58^. High ICAM densities of hundreds of molecules per square micron ensured strong adhesion and is consistent with the expected densities on target cells. Folate receptor antigen densities could be varied over several orders of magnitude, but we focused on high densities of hundreds of molecules per square micron, consistent with the highest potency for these receptors^32^. EC-17 was considered a validated bispecific adapter for this purpose, since it has been shown to image multiple human tumors during cancer surgeries^59^ and because it has also been demonstrated to mediate eradication of solid tumors in animal models^60^.

**Figure 6.**
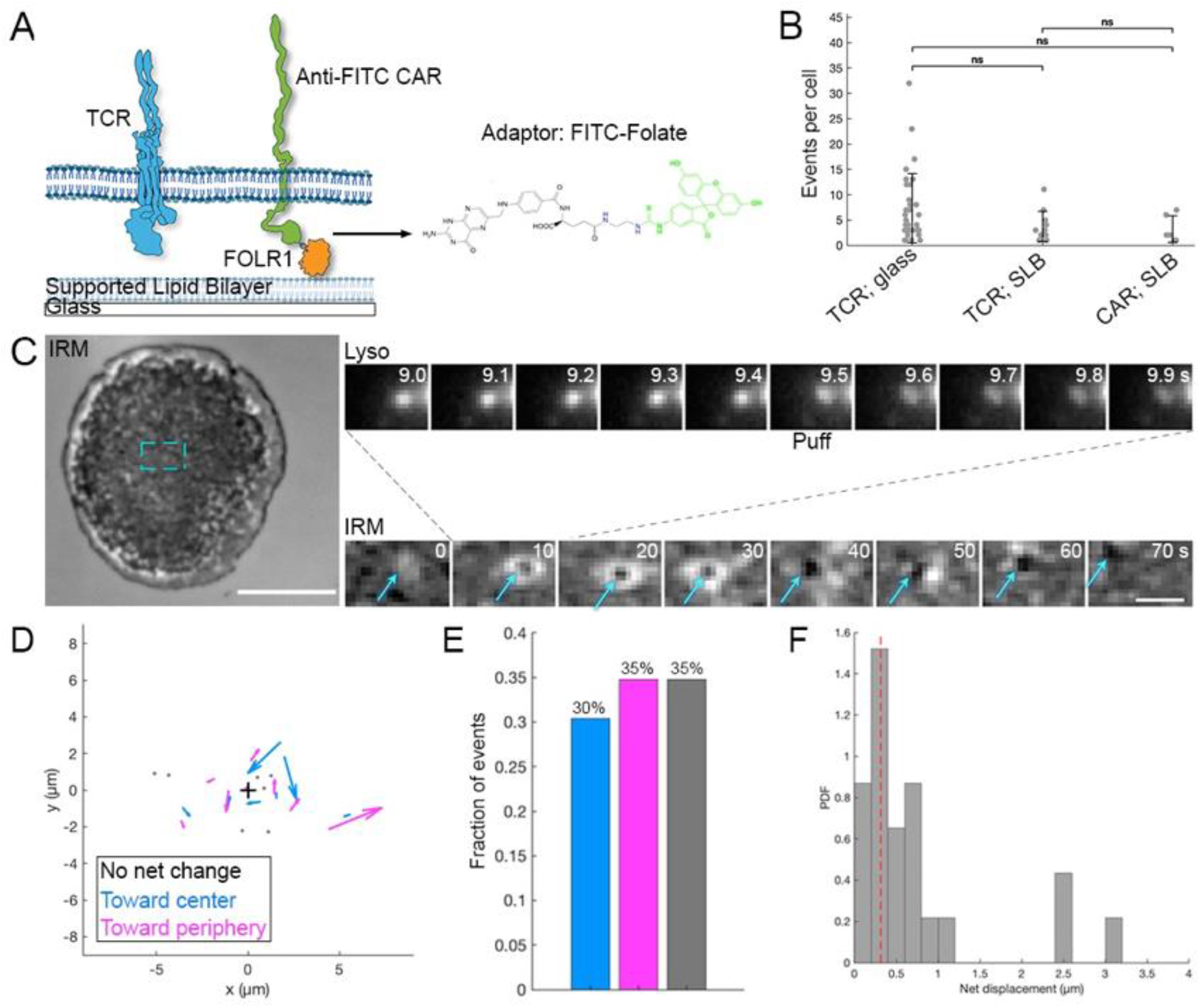
Engineered CAR T cells have similar degranulation patterns on mobile surfaces but have a defect in transporting Mode II particles. **A**. Schematic of engineered T cell stimulation on SLBs. Anti-FITC Chimeric Antigen Receptor (CAR) T cells were prepared in the background of primary human T cells. CAR triggering was achieved using folate receptor alpha (FOLR1) on SLBs and an adaptor molecule FITC-Folate (shown on the right). SLBs presenting a high background of ICAM, similar to other SLBs presenting anti-CD3 (Fig. 5). **B**. Total number of degranulation events per cell on all surfaces tested. ns is not significant. **C**. Representative example of Mode II event on CAR-triggering SLB. Whole cell contact in IRM shown on the left. Scale bar = 5 microns. Cyan box indicates region shown in montages to the right. Top: LysoTracker imaging between sporadic IRM imaging (Bottom montage). In the IRM montage, cyan arrows indicate the Mode II feature. Scale bar = 2 microns. **D**. Displacement vectors for Mode II degranulation events. All events shown across 6 cells where the center of the interface is assigned the coordinate (0,0). **E**. Summary of data in (D) showing the fraction of events for which the Mode II particle moves toward or away from the center, or has no negligible displacement, respectively. **F**. Histogram of Mode II particle displacement. Red, dashed line indicates cutoff for immobile particles (<0.3 micron displacement).

The number of degranulation events per cell (averaging 3-8) were comparable across all substrates and triggering ligands tested (Fig. 6B). The observation of more degranulation events for some T cells on the glass substrate reinforces the strong extent of triggering through mechanical force. Just like Mode II degranulation events on SLBs with TCR triggering, we observed the movement of released particles (Fig. 6C). An example is shown that includes the overall movement of released material toward the center of the contact zone (Fig. 6C; Movie S11). CAR T cells have been shown to form non-classical synapses and characterized by dispersed CAR clustering^11^. We observed similar foci of CAR T cells throughout the interface with some receptors moving toward the periphery and presumably correlated with particle movement (Movie S12). Overall, the frequency of movement toward or away from the center of the junction were equal (Fig. 6D-6E). Additionally, there was an equal probability of no movement, which could also reflect a defect in the reorganization at the interface (Fig. 6D-6E). The overall displacements of released particles were still up to several microns in distance (Fig. 6F). The outward movement of released granules could contribute to the reduced cytotoxic potency in CAR T cells. This would support a model for the native T cell in which mechanical and spatial dependence of granule release at the interface underpins the killing outcome. Together, these results support a model in which the spatiotemporal control of released material add an additional layer of control to T cell cytotoxicity. The insoluble particles may have a longer lifetime and maintain a higher local concentration of cytotoxic molecules that are essential to drive target killing.

## Discussion

The work reported here bridges the molecular mechanisms of T cell degranulation with the mesoscale architecture of the intermembrane interface. This highly regulated space dictates the binding interactions that initiate signal propagation as well as generating the small volume in which cytotoxic molecules accumulate to disrupt and kill target cells. How degranulation is modulated and controlled remains a key, outstanding question. Our combined fluorescence and IRM imaging approach provides an accessible, unified framework for classifying degranulation events as soluble or intact-particle release. Fluorescence imaging enables classification of kinetic features of polarization and degranulation events, while IRM simultaneously reports the presence and behavior of particulate material and T cell membrane topography. The ratio of Mode I:Mode II is regulated by the mobility of the ligands (Fig. 7) and could suggest that changes in tissue stiffness that occur in many disease contexts could regulate cytotoxicity. We also demonstrated that the release of intact particles is accompanied by doming of the overlying membrane (Movie S13) and that the particles can be transported within the interface. Comparison of simulations with IRM sequences put bounds on the physical dimensions of the particles and the local membrane environment (Fig. 7) that are in good agreement with prior reports and add features of the temporal evolution of Mode II degranulation. Collectively, these findings provide a physical insight for understanding how T cells coordinate mechanical and biochemical cues to achieve cytotoxic responses.

**Figure 7.**
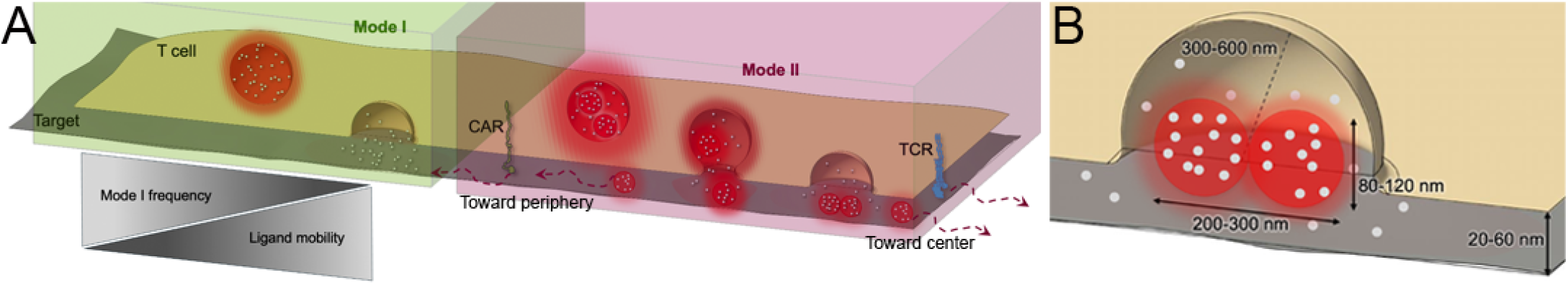
Degranulation is controlled by ligand mobility and released particle transport is regulated by signaling inputs. **A**. Mode I degranulation frequently occurs before Mode II and predominates events on surfaces with immobile ligands. TCR signaling leads to Mode II events whose particles are transported toward the center of the intermembrane junction. CAR signaling can result in the release of Mode II particles that are transported toward the periphery of the junction. **B**. Dimensions of Mode II particles and the membrane landscape based on the findings reported here.

Many mechanistic questions emerge from these results. Primary among these is the licensing of sites for degranulation that, likely, involve signaling, actin rearrangement, assembly of the SNARE-mediated fusion machinery, and membrane choreography. How these components also regulate the number of degranulation events remains a mystery. Many prior studies permitted cells to land prior to spiking cells with calcium and observing a limited number of events^10^. We observed from the onset of signal integration and found that T cells rapidly initiate degranulation, suggesting mechanisms exist to rapidly promote fusion of, primarily, Mode I events. Whether the modes of degranulation and their regulation apply to other immune cells, such as NK cells, is also unclear. Unification of these strategies for profiling both granule contents and membrane geometries with approaches to visualize true cell-cell junctions and mapping to killing outcomes are needed.

Control over the spatiotemporal release of degranulated material is likely a critical determinant of immune protection, tissue homeostasis, and efficacy in immunotherapy. In particular the release of Mode II particles is of great interest given their autonomous killing capacity^5–7^. Concentrating these cytotoxic particles ensures maximal cytotoxicity and minimized collateral damage to neighboring cells. Our results point to another source of off-target toxicity and bystander cell destruction by CAR T cells due to the defects in the temporal regulation of particle localization. Whether different CAR design strategies that combine alternate recognition and signaling domains can modulate the degranulation outcomes, could indicate some of the information processing hierarchy that exists at the level of the receptor itself. Synchronizing input accumulation, polarization, and cytoskeletal rearrangement for degranulation needs to be carefully orchestrated. We predict that Mode II degranulation is delayed ensuring maximal target recognition before the most destructive payloads are released. This temporal control could play a role in antiviral immunity where infected cells could be dispersed and serial killing is required. Thus, a single cytotoxic T cell could deliver a lethal payload and then continue scanning for additional targets. This also distinguishes T cells from immune cells such as mast cells that degranulate in all directions, rather than focus the material release in a directional manner.

Dysregulated degranulation could contribute to tissue damage and autoimmune disease. Secretion of toxic material outside *bona fide* intermembrane junctions would damage to adjacent, healthy cells. How the overlying membrane changes, by gradual collapse or flattening, as two possibilities, could impact the lateral escape of destructive payloads. Especially if the materials released are of the particulate kind, tissue damage could be amplified through a combination of direct cytotoxic damage and stimulation of other immune responses. This could be characteristic of diseases like type I diabetes, multiple sclerosis, and other autoimmune disorders whose hallmarks include chronic cytotoxic T cell activation.

Regulation of degranulation has been of particular interest in antitumor immunity, especially given immunotherapy strategies that mobilize effector responses. Our finding that the ratio of degranulation modes is regulated by mechanics suggests that the physical barriers, altered extracellular matrix (ECM) stiffness, and other immunosuppressive mechanisms in the tumor microenvironment could impair or tune cytotoxic synapses and degranulation capacities. Collectively, these features might also contribute to the substantial lack of success in CAR T cell-based therapies in solid tumors whose biochemical and physical cues are highly complex. Efforts to genetically modify the contents of the particles could also modulate killing outcomes. Thus, efficient mobilization of degranulation, possibly biased toward Mode II, could increase the potency of cytotoxic and CAR T cell function and generate strategies for other cell-based therapies.

## Materials and Methods

### Reagents

Folate Receptor α-10x His (Sino Biological) was conjugated with Alexa Fluor 647 NHS-ester (Fisher Scientific) to achieve a 1:1 protein:dye as previously described^39^. Recombinant ICAM-1-10x His (Acro Biosystems) was used for both coated glass and SLB measurements. Antibody αCD3ε (clone OKT3, BD Biosciences 566685, unconjugated) was used to trigger TCR. Loading of FRα with EC-17 was carried out 12-18 hours prior to imaging and performed fresh for each experiment. EC-17 (AdooQ Bioscience) at 125:1 molar excess was loaded onto FRα-AF647 at 37°C in PBS buffer (1x). Following loading, excess EC-17 was removed by washing.

### T Cell Culture and Lytic Granule Labeling

CAR T cells were prepared by lentiviral transduction in the Low lab according to published protocols and included a 50/50 ratio of CD4+ and CD8+ T cells^57,58^. Primary T cells were purchased as >90% sorted CD8+ T cells (AllCells, Broadview, IL, USA). CAR T cells and non-transduced T cells were cultured in TexMACs GMP phenol-free medium (Miltenyi Biotec) supplemented with 100 IU/mL IL-2 (Miltenyi Biotec) and 2% human serum (Millipore Sigma). Cell passage occurred every 2-3 days to exchange media and maintain cells at a density of 1-3 million cells/mL. The day before imaging, all T cells were exchanged into IL-2 free complete TexMACs medium. Prior to imaging, all T cells were centrifuged (300xg, 4 min, 4°C) and the culture media was exchanged to live cell imaging buffer (LCB; 1 mM CaCl_2_, 2 mM MgCl_2_, 20 mM HEPES, 137 mM NaCl, 5 mM KCl, 0.7 mM Na_2_HPO_4_, 6 mM D-glucose, and 1% m/v BSA) containing LysoTracker DND-99 or LysoTracker DeepRed (Invitrogen) at final 100-200 nM concentration. The cells were incubated at 37°C 5% CO_2_ for 60 minutes. Cells were directly added to coated surfaces or SLBs without additional rinse steps. Remaining cells were stored in an incubator until imaged and used within 2 hours of LysoTracker labeling.

### Bilayer Assembly and Glass Coating

Coated glass surfaces, Hellmanex III (2% v/v in water)-sonicated and rinsed coverslips (#1.5H, 25 mm round, Warner Instruments) were coated with 10 µg/mL poly-L-Lysine (Gibco) diluted in PBS for 10 minutes at room temperature. After multiple rinses with PBS, ICAM-1 or αCD3/ICAM solutions at 10-20 nM ICAM and 10 µg/mL αCD3 as incubated for 60 minutes, covered. After rinsing with PBS, the coverslips were assembled into AttoFluor imaging chambers (ThermoFisher) and media was exchanged to LCB.

Glass-supported lipid bilayer (SLB) membranes were prepared in imaging chambers and functionalized with proteins as previously described^39^. Briefly, SLBs were generated via vesicle fusion. Small unilamellar vesicles (SUVs) were prepared from a lipid mixture consisting of 1,2-dioleoyl-sn-glycero-3-phosphocholine (DOPC, 96 mol%) and 1,2-dioleoyl-sn-glycero-3-[(N-(5-amino-1-carboxypentyl)iminodiacetic acid)succinyl] (nickel salt) (DGS-NTA(Ni), 4 mol%) (Avanti Polar Lipids). Lipids in chloroform were dried under rotovap vacuum (20 mbar) and rotation in 35°C water bath (15 minutes under full vacuum). The lipid film was rehydrated in 18.2 MΩ Milli-Q water (Millipore) to a 1 mg/mL stock. To form SUVs, the lipid suspension was passed 21 times through a 0.2 µm pore polycarbonate filter (Cytiva) using a mini-extruder (Avanti Polar Lipids). SUVs were stored on ice until use that day or freezen (-20 °C) for use within three days.

Cleaned glass coverslips were assembled into imaging chambers. The SUV mixture was diluted to 0.2 mg/mL in 1x PBS, pipetted into the chambers, and mixed via repeated pipetting. Bilayers were allowed to form for 10 minutes at room temperature, followed by extensive rinsing with 1x PBS. To derivatize the surface, bilayers were functionalized with His-tagged recombinant proteins tailored for either T cell receptor (TCR) or chimeric antigen receptor (CAR) triggering. For TCR triggering, SLBs were incubated with Protein L (20 nM) and ICAM-1 (20 nM) followed by αCD3-PE/Dazzle (3.3 nM). For CAR triggering, SLBs were incubated with folate receptor alpha (FRα, 1.4 nM) bound to EC-17, as well as human ICAM-1 (20 nM). All protein incubations were performed for 60 minutes at room temperature with manual mixing. Bilayers were rinsed thoroughly with 1x PBS. Once fully formed and functionalized, SLBs were utilized for imaging within 8 hours of final rinse step.

### Live Cell Imaging

A Nikon Ti2-E Eclipse inverted microscope equipped with a 100x TIRF objective (Nikon), 4 laser lines (405, 488, 561, and 640 nm), Perfect Focus system, motorized TIRF arm, Epifluorescence lamp (X-Cite) arm, and live-cell incubation stage was used for all experiments. A live-cell incubation stage (Tokai Hit) was used to maintain an environment of 37°C and 5% CO_2_ for all live T cell measurements. Excitation used a quad-color excitation cube. An SRIC excitation cube was used for IRM imaging. Data were captured on an EM-CCD (Andro iXon Life). All components, including the motorized stage were controlled by Nikon Elements software. The coated glass or SLB-containing imaging chambers was pre-warmed for 5-10 minutes before imaging. For each chamber, 0.2 million cells were added in a volume of 20-50 µL of LCB.

Two main acquisition sequences were used. To monitor degranulation directly, the LysoTracker channel was streamed in TIRF at 10Hz and interleaved with IRM images every 10 seconds. For imaging to emphasize the membrane geometry, IRM was streamed at 10Hz and interleaved with diascopic or TIRF-based images very 10 seconds.

### ELISA

To assay the ensemble levels of degranulation, T cells were exposed to adhesion only (ICAM-1 only) or stimulating (αCD3/ICAM-1) coated surfaces. These samples were imaged in AttoFluor chambers prior to supernatant capture. After 30 minutes, 800 μL of supernatants were collected and spun down at 1000xg for 10 minutes to remove any cells or debris. The supernatant was pipetted off and stored at -20°C. Supernatants were thawed and an ELISA kit for granzyme B (Invitrogen) was used to measure granzyme B levels.

### Estimating FOLR Ligand Density

The number of FOLR1 molecules on the SLB surface was calculated as described previously^39^ and follows from measuring the integrated fluorescence from one fluorophore-protein conjugate and scaling to the integrated fluorescence over a fixed area.

### IRM Modeling

Mode I parameters: size of dome = 200 nm; h_cell_ (adhesion length) = 45 nm; n_granule fluid_ = 1.36; n_cytoplasm_ = 1.36; n_extracellular medium_ = 1.33; n_glass_ = 1.525; n_lipid_ = 1.486; d_lipid_ = 4; d_water_ = 2.

### Image Analysis

An analysis pipeline, implemented mainly in MATLAB (MathWorks, Natick, MA), was developed (available on GitHub; https://github.com/Low-NamLab) to process raw microscopy data into cell traces in time and corresponding Lytic granule tracking. Preprocessing IRM and DIA channels involved a median image correct for uneven illumination and artifacts (dust, debris) along the optical path. Fluorescence images were EMCCD gain corrected. Binding data for TCR and CAR conditions were shade corrected using calibration images from lipophilic dyes incorporated into planar SLBs (DiO, DiI and DiD, Invitrogen).

Preprocessed data were saved as .TIFF and imported to ImageJ for tracking using the TrackMate plugin^61^. IRM dark spots were tracked manually based on the spot’s contrast and temporal persistence. Once deposited on the surface, these features persisted for the remainder of the acquisition, with contrast varying according to local ruffling of the T cell plasma membrane and the motion of internal organelles.Localizations were connected into tracks using the Linear Assignment Problem (LAP) Tracker with a 0.5 µm linking distance, and gap closing of 1 µm and max 20 frame gap over the track lifetime. Tracks of less than 10 frames in length (1 s real time) were excluded from further analysis. Track coordinates were exported to MATLAB for further analysis.

LysoTracker differs from genetically encoded pH reporters such as pHluorin fusion proteins in that its fluorescence decreases rather than increases upon exocytosis. To classify exocytic events, careful integration of the time-dependent evolution of both granule intensity and radius was conducted. A distinct brightening of signal as the granule fused with the membrane (at a z position in a stronger portion of the TIRF field illumination) followed by the radial dispersion, ‘puff, of that signal followed by quenching. This distinction allows reliable classification of exocytic events without the use of exogenous tagged fusion proteins or additional labeling of lytic granules with lectin markers.

LysoTracker tracking was performed in MATLAB. For LysoTracker, candidate spots were identified using a Laplacian of Gaussian (LoG) filter, with the threshold set to minimize false-positive spots in later frames containing few lytic granules. Candidate spots were then fit with a GPU-accelerated 2D Gaussian algorithm. Spots were linked assuming a maximum diffusivity of 2 µm^2^/s, with a Kalman filter for motion prediction. Tracking was terminated when the fluorescent signal (signal-to-background ratio, SBR > 1) could no longer be well resolved by this algorithm. Tracks shorter than 5 frames (0.5 s real time) were excluded from further analysis.

For classifying events, degranulating lytic granule tracks were first partitioned into time points before and after the puff-annotation frame. The points after the puff were fit with a single exponential, f(x) = A · e^(−Bx)^ + C, where A is the fit amplitude, B is the decay factor (1/τ), and C is a y-offset. Tracks with fewer than three post-puff data points were excluded, as were tracks whose fits returned physically impossible large τ values — for example, dim events whose signal fell below the detection limit before a decay could be reliably measured. Five out of 241 degranulation events were excluded.

## Acknowledgements

We acknowledge the lab of Dr. Philip Low (Purdue University) for providing the primary CAR T cells.

## Notes

### Competing Interest Statement

The authors have declared no competing interest.

## References

1. Metkar, S. S. et al. Cytotoxic Cell Granule-Mediated Apoptosis: Perforin Delivers Granzyme B-Serglycin Complexes into Target Cells without Plasma Membrane Pore Formation. Immunity vol. 16 (2002).

2. Wang, M. S. et al. Mechanically active integrins target lytic secretion at the immune synapse to facilitate cellular cytotoxicity. Nat. Commun. 13, 1–15 (2022).

3. Pui-Yan Ma, V. et al. The magnitude of LFA-1/ICAM-1 forces fine-tune TCR-triggered T cell activation. Sci. Adv. 8, 1–17 (2022).

4. Cassioli, C. & Baldari, C. T. The Expanding Arsenal of Cytotoxic T Cells. Front. Immunol. 13, 1–7 (2022).

5. Chang, H. F. et al. Identification of distinct cytotoxic granules as the origin of supramolecular attack particles in T lymphocytes. Nat. Commun. 13, 1–15 (2022).

6. Bálint et al. Supramolecular attack particles are autonomous killing entities released from cytotoxic T cells. Science (1979). 368, 897–901 (2020).

7. Cassioli, C. et al. Activation-induced thrombospondin-4 works with thrombospondin-1 to build cytotoxic supramolecular attack particles. Proc. Natl. Acad. Sci. U. S. A. 122, (2025).

8. Ming, M., Schirra, C., Becherer, U., Stevens, D. R. & Rettig, J. Behavior and properties of mature lytic granules at the immunological synapse of human cytotoxic T lymphocytes. PLoS One 10, (2015).

9. Chang, H. F. et al. Cytotoxic granule endocytosis depends on the Flower protein. Journal of Cell Biology 217, 667–683 (2018).

10. Chitirala, P. et al. Studying the biology of cytotoxic T lymphocytes in vivo with a fluorescent granzyme B-mTFP knock-in mouse. Elife 9, 1–19 (2020).

11. Davenport, A. J. et al. Chimeric antigen receptor T cells form nonclassical and potent immune synapses driving rapid cytotoxicity. Proceedings of the National Academy of Sciences 115, E2068–E2076 (2018).

12. Stinchcombe, J. C. et al. Ectocytosis renders T cell receptor signaling self-limiting at the immune synapse. Science (1979). 380, 818–823 (2023).

13. Jenkins, M. R., Tsun, A., Stinchcombe, J. C. & Griffiths, G. M. The Strength of T Cell Receptor Signal Controls the Polarization of Cytotoxic Machinery to the Immunological Synapse. Immunity 31, 621–631 (2009).

14. Pattu, V. et al. Syntaxin7 is required for lytic granule release from cytotoxic T lymphocytes. Traffic 12, 890–901 (2011).

15. Qu, B. et al. Docking of Lytic Granules at the Immunological Synapse in Human CTL Requires Vti1b-Dependent Pairing with CD3 Endosomes. The Journal of Immunology 186, 6894–6904 (2011).

16. Quintana, A. et al. T Cell Activation Requires Mitochondrial Translocation to the Immunological Synapse. 10.1073pnas.0703126104 (2007).

17. Chang, H. F. et al. Preparing the lethal hit: interplay between exo- and endocytic pathways in cytotoxic T lymphocytes. Cellular and Molecular Life Sciences vol. 74 399–408 Preprint at 10.1007/s00018-016-2350-7 (2017).

18. Martina, J. A. et al. Imaging of lytic granule exocytosis in CD8+ cytotoxic T lymphocytes reveals a modified form of full fusion. Cell. Immunol. 271, 267–279 (2011).

19. McAffee, D. B. et al. Discrete LAT condensates encode antigen information from single pMHC:TCR binding events. Nat. Commun. 13, (2022).

20. Lin, J. J. Y. et al. Mapping the stochastic sequence of individual ligand-receptor binding events to cellular activation: T cells act on the rare events. Sci. Signal. 12, (2019).

21. O’Donoghue, G. P., Pielak, R. M., Smoligovets, A. A., Lin, J. J. & Groves, J. T. Direct single molecule measurement of TCR triggering by agonist pMHC in living primary T cells. Elife 2013, 1–16 (2013).

22. Manz, B. N., Jackson, B. L., Petit, R. S., Dustin, M. L. & Groves, J. T-cell triggering thresholds are modulated by the number of antigen within individual T-cell receptor clusters. Proc. Natl. Acad. Sci. U. S. A. 108, 9089–9094 (2011).

23. Purbhoo, M. A., Irvine, D. J., Huppa, J. B. & Davis, M. M. T cell killing does not require the formation of a stable mature immunological synapse. Nat. Immunol. 5, 524–530 (2004).

24. Irvine, D. J., Purbhoo, M. A., Krogsgaard, M. & Davis, M. M. Direct observation of ligand recognition by T cells. Nature 419, 845–849 (2002).

25. Patel, K. K., Tariveranmoshabad, M., Kadu, S., Shobaki, N. & June, C. From concept to cure: The evolution of CAR-T cell therapy. Molecular Therapy 33, 2123–2140 (2025).

26. Ramello, M. C. et al. An immunoproteomic approach to characterize the CAR interactome and signalosome. Sci. Signal. 12, (2019).

27. Salter, A. I. et al. Comparative analysis of TCR and CAR signaling informs CAR designs with superior antigen sensitivity and in vivo function. Sci. Signal. 14, (2021).

28. Dong, R. et al. Rewired signaling network in T cells expressing the chimeric antigen receptor ( CAR ) . EMBO J. 39, 1–14 (2020).

29. Stinchcombe, J. C., Majorovits, E., Bossi, G., Fuller, S. & Griffiths, G. M. Centrosome polarization delivers secretory granules to the immunological synapse. Nature 443, 462–465 (2006).

30. Dillard, P., Varma, R., Sengupta, K. & Limozin, L. Ligand-Mediated Friction Determines Morphodynamics of Spreading T Cells. Biophys. J. 107, 2629–2638 (2014).

31. Limozin, L. & Sengupta, K. Modulation of Vesicle Adhesion and Spreading Kinetics by Hyaluronan Cushions. Biophys. J. 93, 3300–3313 (2007).

32. Gudipati, V. et al. Inefficient CAR-proximal signaling blunts antigen sensitivity. Nat. Immunol. 21, 848–856 (2020).

33. Gwalani, L. A. & Orange, J. S. Single Degranulations in NK Cells Can Mediate Target Cell Killing. The Journal of Immunology 200, 3231–3243 (2018).

34. Monzel, C. & Sengupta, K. Measuring shape fluctuations in biological membranes. J. Phys. D Appl. Phys. 49, (2016).

35. Limozin, L. & Sengupta, K. Quantitative reflection interference contrast microscopy (RICM) in soft matter and cell adhesion. ChemPhysChem 10, 2752–2768 (2009).

36. Estl, M. et al. Various stages of immune synapse formation are differently dependent on the strength of the TCR stimulus. Int. J. Mol. Sci. 21, (2020).

37. Melenhorst, J. J. et al. Decade-long leukaemia remissions with persistence of CD4+ CAR T cells. Nature 602, 503–509 (2022).

38. Anikeeva, N. et al. Efficient killing of tumor cells by CAR-T cells requires greater number of engaged CARs than TCRs. Journal of Biological Chemistry 297, (2021).

39. Scrudders, K. L. et al. CAR T Cell Cytotoxic Responses Are Rapidly Generated and Sensitive to the Unligated TCR.

40. Govendir, M. A. et al. T cell cytoskeletal forces shape synapse topography for targeted lysis via membrane curvature bias of perforin. Dev. Cell 57, 2237-2247.e8 (2022).

41. Lemaître, F. et al. Unveiling the molecular architecture of T cells and immune synapses with cryo-expansion microscopy. Cell Rep. 45, (2026).

42. Llobet, A., Beaumont, V. & Lagnado, L. Neurotechnique Real-Time Measurement of Exocytosis and Endocytosis Using Interference of Light. Neuron vol. 40 http://www.neuron.org/cgi/content/full/40/ (2003).

43. Liu, D., Martina, J. A., Wu, X. S., Hammer, J. A. & Long, E. O. Two modes of lytic granule fusion during degranulation by natural killer cells. Immunol. Cell Biol. 89, 728–738 (2011).

44. Wiegand, G., Neumaier, K. R. & Sackmann, E. Microinterferometry: three-dimensional reconstruction of surface microtopography for thin-film and wetting studies by reflection interference contrast microscopy (RICM). Appl. Opt. 37, 6892 (1998).

45. Rädler, J. & Sackmann, E. Imaging optical thicknesses and separation distances of phospholipid vesicles at solid surfaces. Journal de Physique II 3, 727–748 (1993).

46. Abdelrahman, A., Smith, A. S. & Sengupta, K. Observing Membrane and Cell Adhesion via Reflection Interference Contrast Microscopy. in Methods in Molecular Biology vol. 2654 123–135 (Humana Press Inc., 2023).

47. Robert, P., Sengupta, K., Puech, P.-H., Bongrand, P. & Limozin, L. Tuning the Formation and Rupture of Single Ligand-Receptor Bonds by Hyaluronan-Induced Repulsion. Biophys. J. 95, 3999–4012 (2008).

48. Taylor, R. W. & Sandoghdar, V. Interferometric Scattering Microscopy: Seeing Single Nanoparticles and Molecules via Rayleigh Scattering. Nano Lett. 19, 4827–4835 (2019).

49. Shaw, A. S. & Dustin, M. L. Making the T Cell Receptor Go the Distance: Review A Topological View of T Cell Activation. Immunity vol. 6 (1997).

50. Leithner, A. et al. Mixed-mobility supported lipid bilayers uncover the role of immobilized ICAM1 on T cell activation and immune synapse organization. Proceedings of the National Academy of Sciences 123, (2026).

51. Mossman, K. D., Campi, G., Groves, J. T. & Dustin, M. L. Immunology: Altered TCR signaling from geometrically repatterned immunological synapses. Science (1979). 310, 1191–1193 (2005).

52. Groves, J. T. & Dustin, M. L. Supported planar bilayers in studies on immune cell adhesion and communication. Journal of Immunological Methods vol. 278 19–32 Preprint at 10.1016/S0022-1759(03)00193-5 (2003).

53. Grakoui, A. et al. The immunological synapse: A molecular machine controlling T cell activation. Science (1979). 285, 221–227 (1999).

54. Dustin, M. L. The immunological synapse. Cancer Immunol. Res. 2, 1023–1033 (2014).

55. DeMond, A. L., Mossman, K. D., Starr, T., Dustin, M. L. & Groves, J. T. T cell receptor microcluster transport through molecular mazes reveals mechanism of translocation. Biophys. J. 94, 3286–3292 (2008).

56. Barden, M. et al. CAR and TCR form individual signaling synapses and do not cross-activate, however, can co-operate in T cell activation. Front. Immunol. 14, (2023).

57. Lu, Y. J. et al. Preclinical evaluation of bispecific adaptor molecule controlled folate receptor CAR-T cell therapy with special focus on pediatric malignancies. Front. Oncol. 9, 1–20 (2019).

58. Lee, Y. G. et al. Regulation of CAR T cell-mediated cytokine release syndrome-like toxicity using low molecular weight adapters. Nat. Commun. 10, 1–11 (2019).

59. Low, P. S. & Kularatne, S. A. Folate-targeted therapeutic and imaging agents for cancer. Current Opinion in Chemical Biology vol. 13 256–262 Preprint at 10.1016/j.cbpa.2009.03.022 (2009).

60. Lu, Y. & Low, P. S. Folate targeting of haptens to cancer cell surfaces mediates immunotherapy of syngeneic murine tumors. Cancer Immunology, Immunotherapy 51, 153–162 (2002).

61. Ershov, D. et al. TrackMate 7: integrating state-of-the-art segmentation algorithms into tracking pipelines. Nat. Methods 19, 829–832 (2022).

