## Supplemental Information for "T cell degranulation and payload delivery are modulated in space and time"

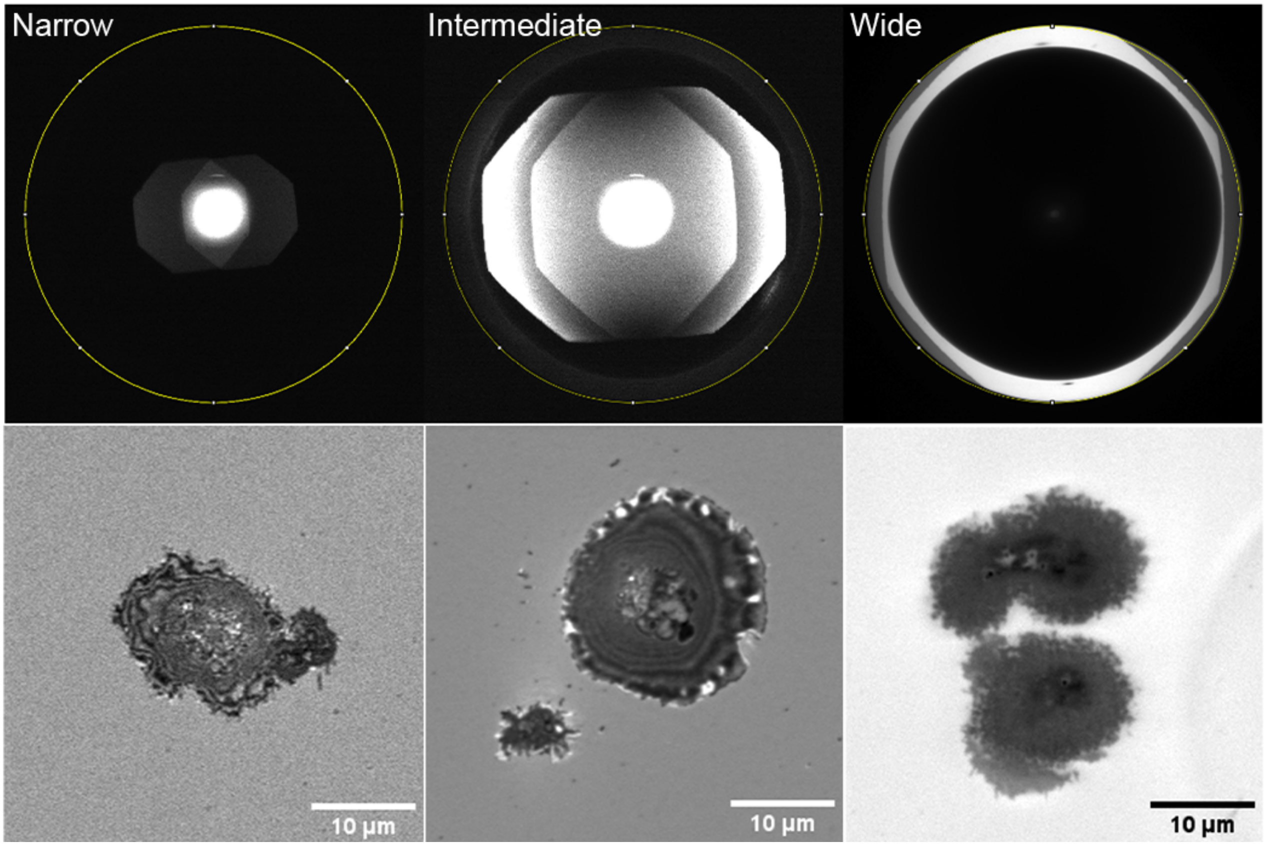


**Figure S1. IRM aperture setting.** Aperture diaphragm imaged on a back aperture camera (top row) showing narrow, intermediate, and wide positions, respectively. Below each image is a corresponding image of T cells on a glass substrate at the corresponding position.

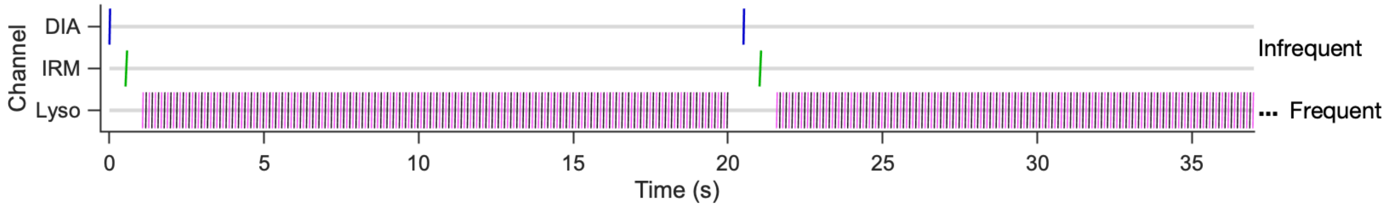


**Figure S2. Multichannel acquisition scheme for degranulation imaging.** Interleaved protocol using streaming of frequent LysoTracker (640 nm) TIRF imaging (10 Hz) and infrequent imaging of each of diascopic (DIA) and IRM (every 20 s), separated by filter switches. Channels are shown on the y axis.

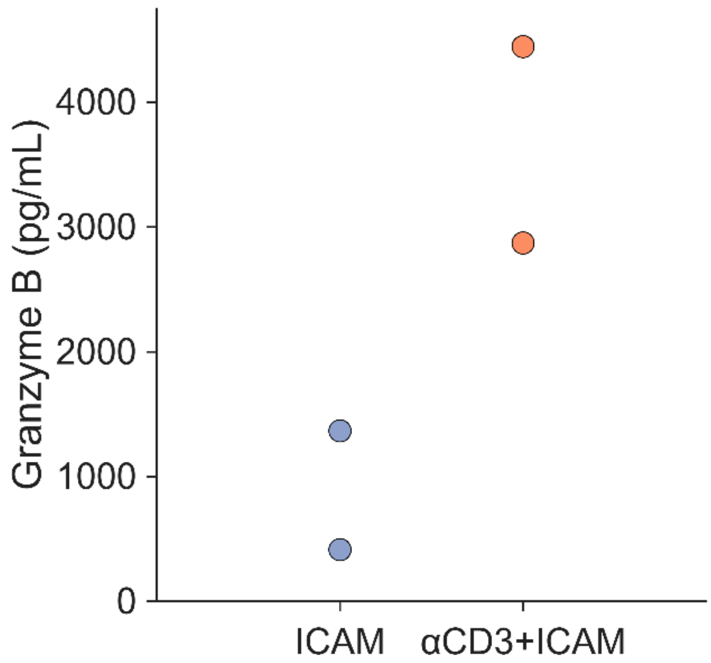


**Figure S3. Total T cell degranulation on coated glass substrates.** Granzyme B release measured by ELISA from primary human T cells on coated glass. Coverslips were coated with poly-L-lysine (1 μg/mL) followed by either ICAM alone (20 nM) or anti-CD3 (10 μg/mL) plus ICAM (20 nM). Cells were imaged on coated surfaces at 37°C for up to 30 minutes, after which supernatants were collected and assayed for granzyme B. Granzyme B levels were elevated on the activating (anti-CD3 + ICAM) surfaces, consistent with robust degranulation. Data from 2 technical replicates.

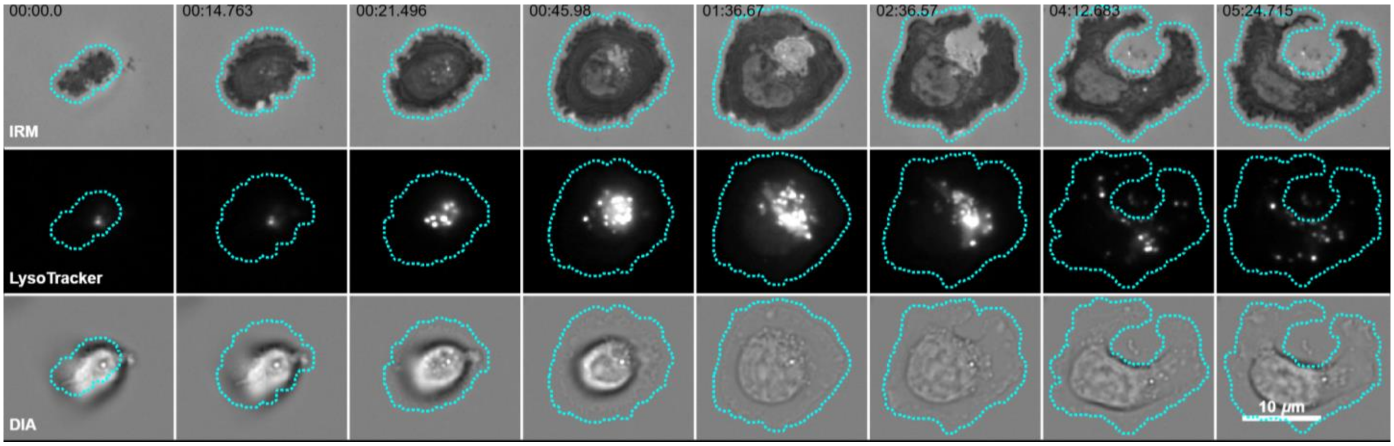


**Figure S4. Complete T cell retraction montages, corresponding to Figure 2C.** Primary CD8+ T cell degranulation on anti-CD3 + ICAM-coated glass. Cell undergoes retraction, exposing Mode II-released particles in all channels. Scale bar = 10 microns.

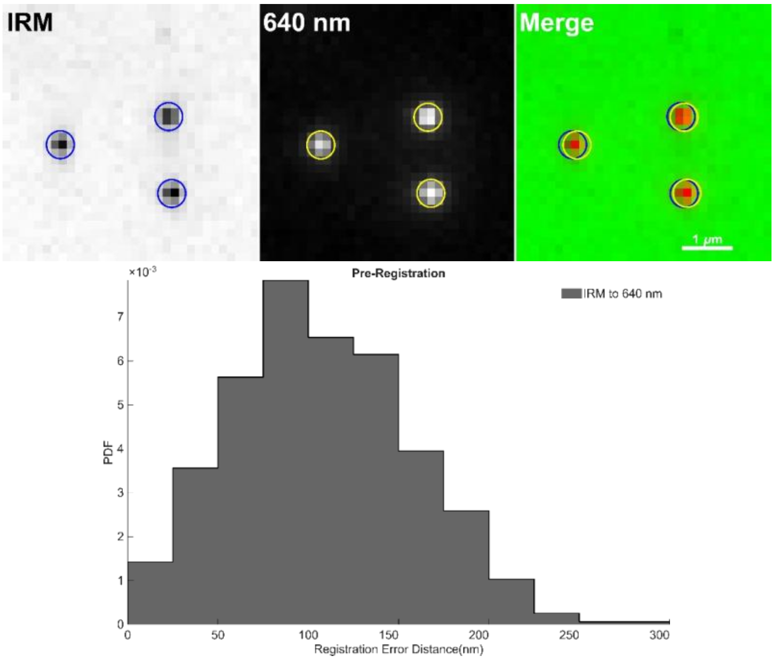


**Figure S5. IRM and TIRF channel registration.** Registration of IRM and TIRF (640 nm) channels was verified using 100 nm TetraSpeck beads deposited on poly-L-lysine-coated glass. Median-filtered IRM objects were localized and compared to coordinates from fitting the fluorescent beads. (Top). The IRM and 640 nm channels achieved sub-pixel registration. No additional registration transformation was applied to raw data (Bottom).

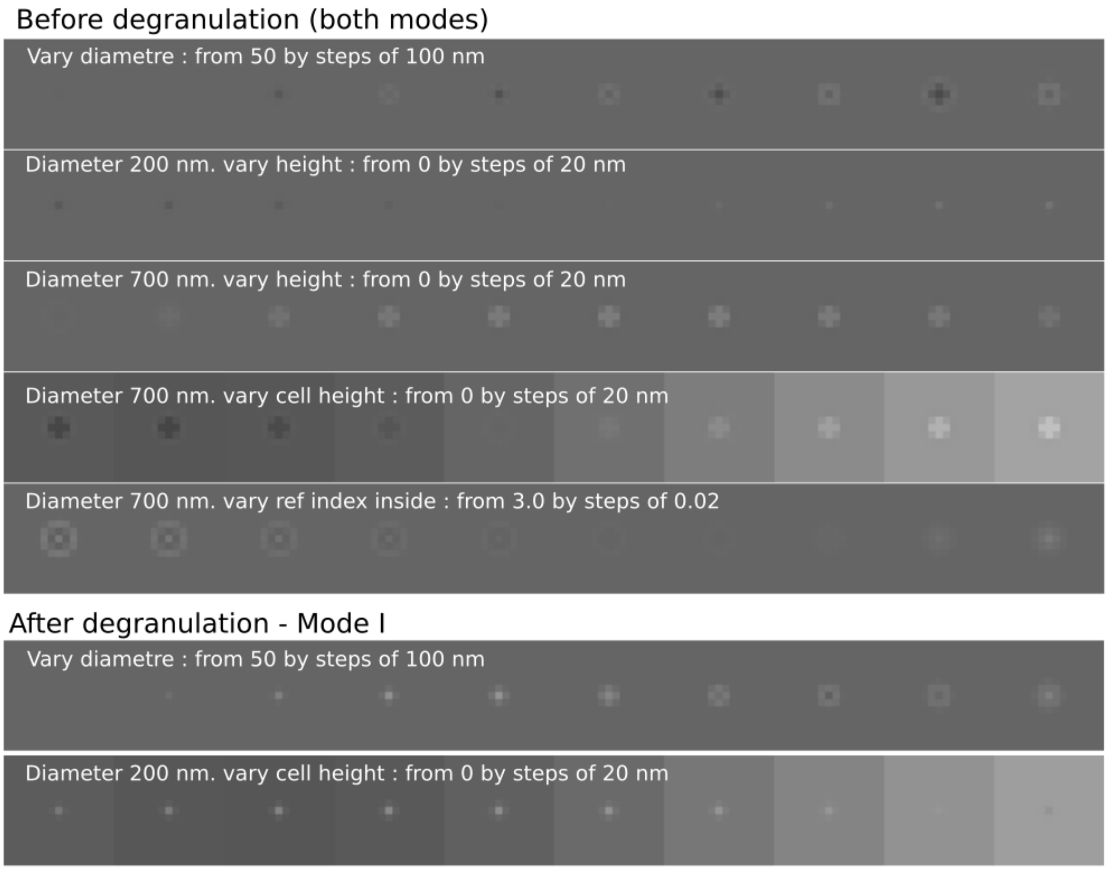


**Figure S6. Complete parameter scanning for Mode I and Mode II events**. Simulated IRM signatures for Mode I and Mode II events before degranulation (above) and Mode I events after degranulation (below).

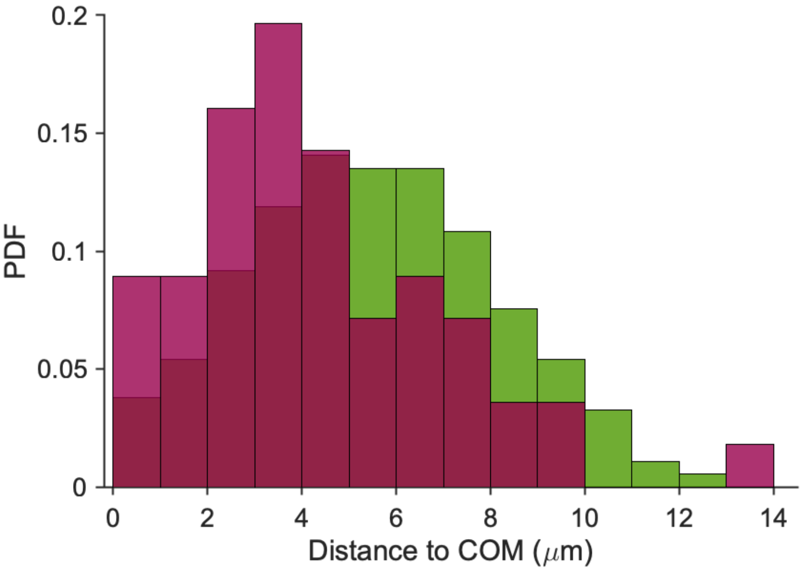


**Figure S7. Mode I events occur more toward the periphery of the interface.** Histogram of Mode I (green) and Mode II events puff locations relative to the center of mass (COM) for that cell.

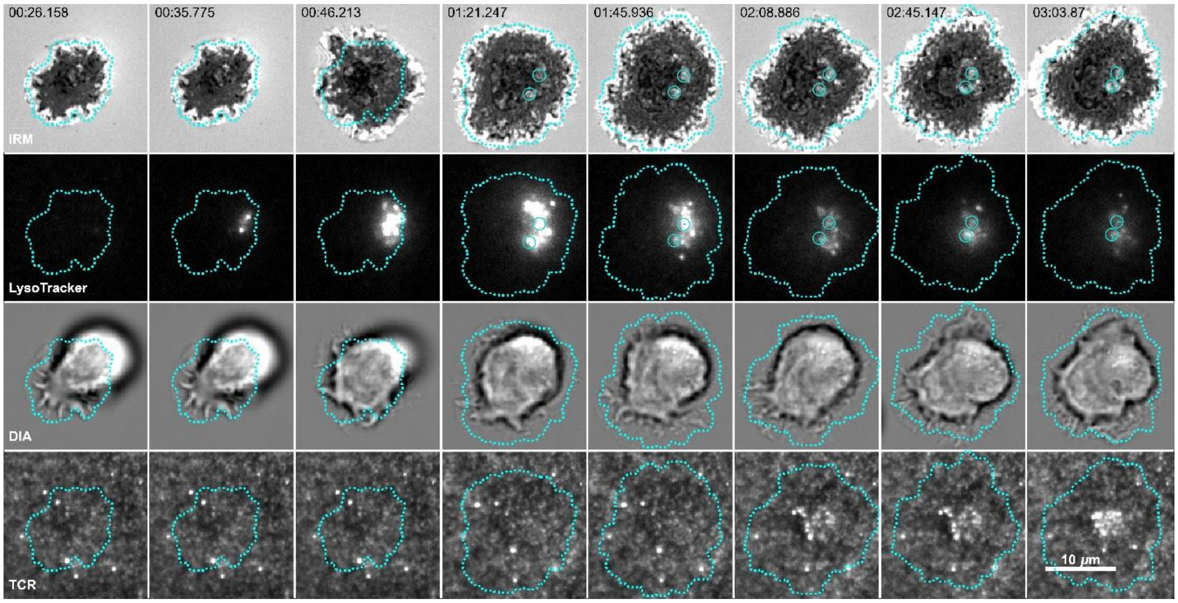


**Figure S8. Released particles centralize following degranulation on anti-CD3 SLBs.** Example montage of a released lytic particle (Mode II) moving toward the center of the interface after release on an anti-CD3 SLB. Centralization coincides with TCR-triggering-induced retrograde flow toward the cell center. Scale bar = 10 microns.

**Supplemental Methods**

IRM simulations were undertaken using original code written in Python. An IRM image was generated via interference of light from reflecting layers of the experimental sample, labeled with an index (i) ranging from 0 to n, each characterized by a refractive index (n_i) and a thickness (d_i; set to infinity for the first and last layers corresponding to the glass support and cell cytoplasm, respectively). The incident monochromatic light (wavelength 560 nm) was approximated to be perpendicular to the layers. The granule objects have radial symmetry and, therefore, simulations considered the shape along a radial line generated from simple geometrical constructions. At each point x along this line, the n_i and d_i for all the layers in the light path were generated. Simulations were performed at 1 nm lateral resolution. Standard RICM equations[35] were applied to obtain the final IRM profile. The profile was rotated around its center to generate the final radially symmetric image. The parameters used to simulate each category of objects are shown in the table below.

| | **Parameter** | **Symbol** | **Value (unless explicitly varied)** |
|---|---|---|---|
| **Cell body** | | | |
| | Cytoplasmic refractive index | n_cell | 1.36 |
| | Height of the cell membrane | h_adh | 45 nm |
| **Before degranulation: Mode I or II** | | | |
| | Diameter of granule | r_sphere | 200 nm/800nm |
| | Height of the bottom of granule | hCntr | 0 |
| | Refractive index of the contents | n_in | 1.36 |
| **After degranulation Mode I** | | | |
| | Diameter of membrane dome | r_sphere | 200 nm |
| **After degranulation Mode II** | | | |
| | Diameter of membrane dome | r_sphere | 800 nm |
| | Diameter and height of deposited particle | d_SMAP/h_SMAP | 200 nm/100 nm |
| | Refractive index of deposited particle | n_SMAP | 1.4 |
| **Fixed Material Parameters** | | | |
| | Refractive index of outer medium | n_out | 1.333 |
| | Refractive index of glass | n_glass | 1.525 |
| | Refractive index of membrane | n_lipid | 1.486 |
| | Thickness of membrane | d_lipid | 4 nm |

To compare the simulated images to the normalized experimental images, the background, corresponding to reflection from glass, was set to 1 and the rest of the image was normalized, accordingly, for each case. The simulated images were Gaussian blurred (sigma=50) and binned to achieve a pixel size of 160 nm, corresponding to experimental camera pixel size. The simulation size was chosen so that after binning, the size of the simulated image matched the size of the experimental image.

Simulations were run for chosen parameters. Each run output 3 important stacks – the stack of simulated images with 1 nm lateral pixel size, the blurred image stack to be compared with experiments, and a representation of the layer geometries. The stacks were exported as tif-stacks, visualized in ImageJ, and finally presented as montages with grey-scale contrast set to 0 to 2.

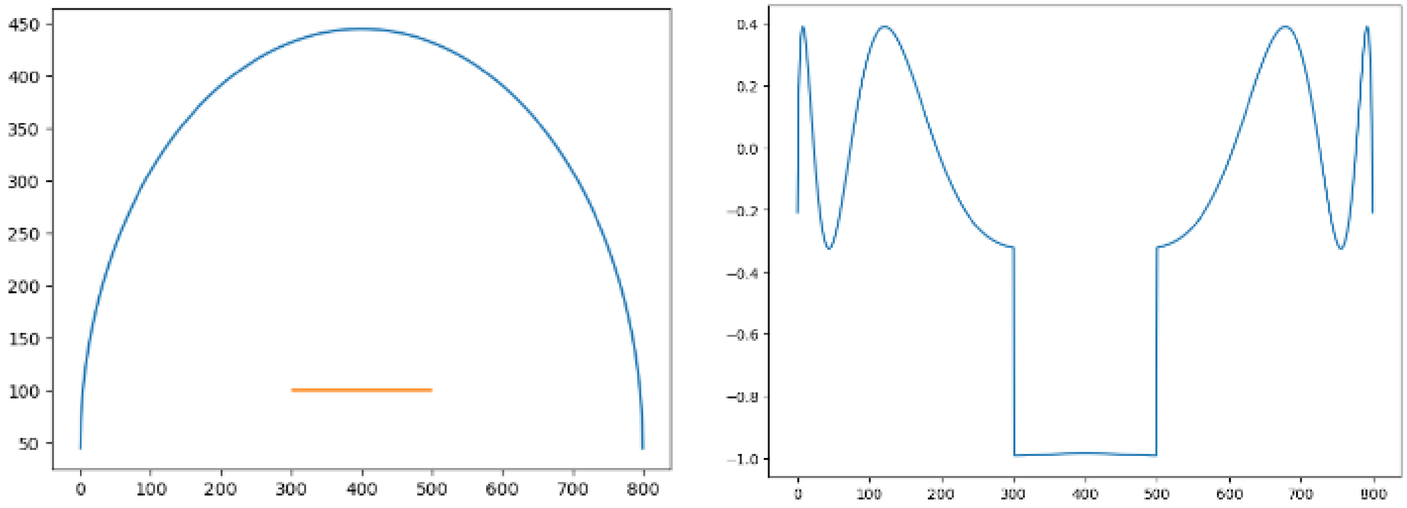


Example output (above) - layer geometry and IRM line profile for the case: after degranulation Mode II. The orange line shows the deposited particle and the blue dome depicts the membrane (Left). The corresponding IRM profile shows a dip in the middle corresponding to the presence of the particle; and fringes corresponding to the domed membrane shape (right). All axis labels are in nm, except the vertical axes for IRM profile. Note that glass background is set to zero.

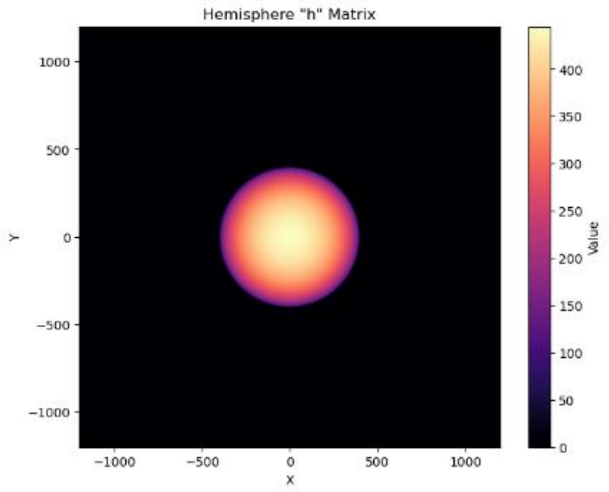


Generated 3D shape of the dome depicted using a color code with scale in nm. x-y scales are in pixel – each 1 nm x 1 nm. Image was generated by rotating the dome profile shown above.

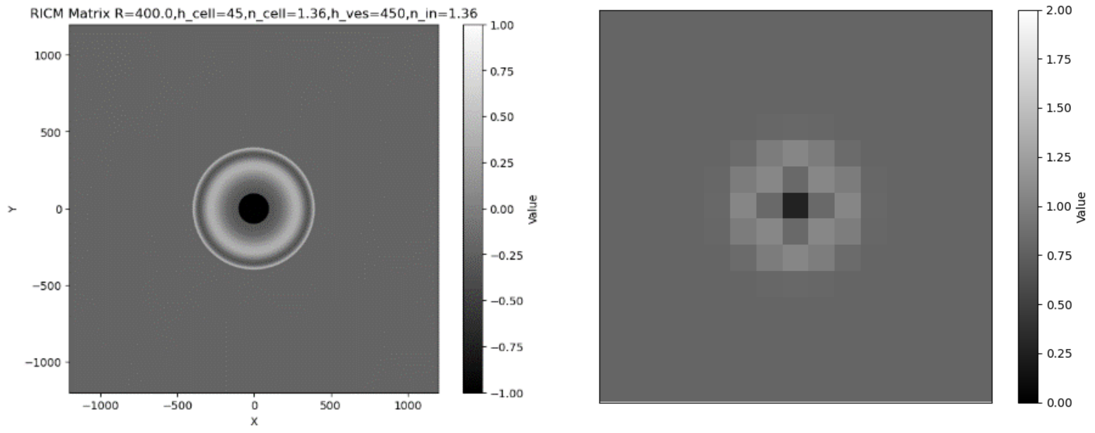


IRM image generated by rotating the IRM profile reported above (left), and the final IRM image for comparison to experiments produced after appropriate normalization, blurring and binning (right). X- and y-scales in nm. Data displayed using normalized intensity.

**Supplemental Movies**

**Movie 1. Primary T cell contact, spreading and lytic granule polarization.** Primary CD8+ T cells loaded with LysoTracker Deep Red were imaged landing on a glass coverslip coated with poly-D-lysine, anti-CD3, and ICAM-1. Cells were imaged using rapid interference reflection microscopy (IRM) acquired at ~8 Hz, interleaved with diascopic and LysoTracker TIRF-based channels every 16 seconds. Here, a T cell undergoes contact, spreading, and lytic granule polarization. Later, membrane retraction exposes released lytic granule particles (Mode 2) that were under the cell. Acquired at ~8 Hz and played at 10x speed. Scale bar = 3 microns.

**Movie 2. Example Mode 1 degranulation event characterized by rapid fluorescence decay.** Primary CD8+ T cells loaded with LysoTracker Deep Red interacting with a poly-D-lysine, anti-CD3, and ICAM-1-coated glass surface. Zoom of an isolated Mode 1 degranulation event, identified by the radial dispersion or "puff" of the LysoTracker signal, followed by a characteristic rapid fluorescent decay and signal loss. Data are shown from 4 seconds preceding to 4 seconds after the degranulation event. The corresponding integrated intensity in photons is derived from a 2D Gaussian fit of the spot as a function of time. Acquired at ~10 fps and played at 2.1x speed (slowed during the puff). Scale bar = 1 micron.

**Movie 3. Example Mode 2 degranulation event characterized by particle release and prolonged fluorescence decay.** Primary CD8+ T cells loaded with LysoTracker Deep Red interacting with a poly-D-lysine, anti-CD3, and ICAM-1-coated glass surface. Zoom of an isolated Mode II degranulation event, identified by prolonged fluorescence decay due to particle release after the degranulation puff. Acquired at ~10 Hz and played at 2.1x speed (slowed during the puff). Scale bar = 1 micron.

**Movie 4. Simulation of Mode I degranulation, before release, under modulation of distance to surface.** Parameter scanning of Mode I degranulation with decreasing distance to the plasma membrane in steps of 50 nm from 450 nm above the surface. Simulated image (left) and corresponding input geometry (right). Scale bar = 200 nm.

**Movie 5. Simulation of Mode II degranulation, before release, under modulation of distance to surface.** Parameter scanning of Mode II degranulation with decreasing distance to the plasma membrane in steps of 50 nm from 800 nm above the surface. Simulated image (left) and corresponding input geometry (right). Scale bar = 200 nm.

**Movie 6. Simulation of Mode I degranulation under modulation of dome dimensions.** Parameter scanning of Mode I degranulation with dome diameter decreasing in steps of 50 nm from 450 nm. Simulated image (left) and corresponding input geometry (right). Scale bar = 200 nm.

**Movie 7. Simulation of Mode II degranulation under modulation of dome dimensions.** Parameter scanning of Mode II degranulation with dome diameter decreasing in steps of 50 nm from 800 nm. Simulated image (left) and corresponding input geometry (right). Scale bar = 200 nm.

**Movie 8. Simulation of Mode II degranulation under modulation of overlying membrane geometry.** Parameter scanning of Mode II degranulation with dome diameter, assuming a spherical geometry, decreasing in steps of 50 nm from 1000 nm. Simulated image (left) and corresponding input geometry (right). Scale bar = 200 nm.

**Movie 9. Simulation of Mode II degranulation under modulation of overlying membrane geometry.** Parameter scanning of Mode II degranulation with dome diameter, assuming the membrane flattens, decreasing in steps of 50 nm from 1000 nm. Simulated image (left) and corresponding input geometry (right). Scale bar = 200 nm.

**Movie 10. T cell degranulation and mature immunological synapse formation on a supported lipid bilayer displaying mobile adhesion and TCR ligands.** Primary CD8+ T cells landing on a supported lipid bilayer (SLB) derivatized with ICAM-1 and anti-CD3 antibody molecules. LysoTracker signal was imaged rapidly using TIRF at ~10 Hz, while the IRM and TCR binding (anti-CD3) channels were acquired at slower intervals (every 10 seconds). Annotations indicate degranulation events (white arrows) and the formation of a mature, immunological synapse where the anti-CD3 antibodies aggregate in the center of the contact zone. Acquired at ~10 Hz and shown at 11x speed. Scale bar = 3 microns.

**Movie 11. Single particle tracking of a Mode 2 degranulation event by a CAR T cell that forms a typical synapse.** Primary T cell expressing chimeric antigen receptor (CAR) landing on an SLB presenting ICAM and the antigen FOLR1. A Mode 2 degranulation event, monitored by rapid LysoTracker imaging occurs and is annotated. The LysoTracker channel was acquired at 2 Hz, while CAR binding and IRM channels were acquired every 20 seconds. Released particles are identified in the IRM channel by a white circle and tracked with a yellow tail that fades every eight IRM frames. Particle remains toward the center of the interface. Acquired at 2 Hz and shown at 9.5x speed. Scale bar = 3 microns.

**Movie 12. Single particle tracking of a Mode 2 degranulation event by a CAR T cell that forms a noncanonical synapse.** Primary T cell expressing chimeric antigen receptor (CAR) landing on an SLB presenting ICAM and the antigen FOLR1. A Mode 2 degranulation event occurs and is annotated. The LysoTracker channel was acquired at ~8 Hz, while IRM and CAR binding channels were acquired every 10 seconds. Released particles are identified in the IRM channel by a white circle and tracked with a yellow tail that fades every eight IRM frames. Particle is transported toward the periphery of the interface. Acquired at ~8 Hz and shown at 9x speed. Scale bar = 3 microns.

**Movie 13. Plasma membrane doming following T cell degranulation.** Primary CD8+ T cells loaded with LysoTracker Deep Red interacting with a poly-D-lysine, anti-CD3, and ICAM-1-coated glass surface. Multiple Mode 2 degranulation events are annotated. Notably, one event is highlighted to be followed by substantive, micron-scale doming of the overlying T cell plasma membrane, lifting away from the glass surface. Both LysoTracker and IRM channels are shown. Acquired at ~8 fps and shown at 1.5x speed. Scale bar = 3 microns.
